# HTT is a repressor of ABL activity required for APP induced axonal growth

**DOI:** 10.1101/679381

**Authors:** Claire Marquilly, Germain Busto, Brittany S. Leger, Edward Giniger, James A. Walker, Lee G. Fradkin, Jean-Maurice Dura

**Affiliations:** IGH, Centre National de la Recherche Scientifique, Univ Montpellier, Montpellier, France; McGill University Centre for Research in Neuroscience, Department of Neurology and Neurosurgery, Research Institute of the McGill University Health Centre, Montreal, QC, Canada; Center for Genomic Medicine, Massachusetts General Hospital, Boston, MA 02114, USA; Intramural Research Program, NINDS, NIH, Bethesda, MD 20892, USA; Department of Neurology, Massachusetts General Hospital, Harvard Medical School, Boston, USA; Cancer Program, Broad Institute of MIT and Harvard, Cambridge, MA 02142, USA; Department of Neurobiology, University of Massachusetts Medical School, 364 Plantation Ave. Worcester, MA 01655, USA

**Keywords:** HTT, ABL, FRET, axonal growth, Appl signaling, mushroom body

## Abstract

ABL tyrosine kinase activity controls several aspects of development including axon patterning. Amyloid precursor protein (APP) is linked to Alzheimer’s disease and previous work established that ABL is a downstream effector in an *Appl*, the *Drosophila* App ortholog, signaling pathway which modulates axon outgrowth in the mushroom bodies (MBs), the fly memory center. Here we show that *Abl* is required for the MB neuron axonal growth. Importantly, both *Abl* overexpression and lack of expression produce a similar phenotype in the MBs indicating the necessity of tightly regulating ABL activity. We find that the fly huntingtin protein (HTT), the homolog of the protein involved in Huntington’s disease, behaves genetically as a repressor of ABL activity. Supporting this, FRET-based measurements of *in vivo* ABL activity in the MBs reveal a clear increase in its activity when HTT levels are reduced. Thus, in addition to its many other reported roles, HTT acts as a negative regulator of ABL activity, at least in the MBs, to maintain its appropriate physiological levels necessary for axon growth.

## INTRODUCTION

Tumorigenesis and neurodegeneration may be two sides of the same coin. Indeed, defining the overlap of molecular pathways implicated in cancer and neurodegeneration may open the door to novel therapeutic approaches for both groups of disorders (Staropoli, 2008). Interestingly, it has been proposed that, in the non-pathologic physiological situation, the neurodegenerative Amyloid precursor protein (APP) recruits the oncogenic Abelson (Abl) kinase in order to promote axonal outgrowth (Soldano et al., 2013). Correlative studies have highlighted a decreased cancer incidence in the neurodegenerative disorder Huntington’s disease (HD) population and both wild-type and mutant huntingtin (HTT) have been implicated in tumor progression (Thion and Humbert, 2018). It is, therefore, tempting to propose that different neurodegenerative diseases (ND) may share components and mechanisms and that Abl may also have a role in ND.

The Abl-family of non-receptor tyrosine kinase includes human ABL1 and ABL2 as well as the *Drosophila Abl*, all of which share a conserved domain structure along much of the protein. Each ABL protein contains an SH3-SH2-TK (Src homology 3-Src homology 2-tyrosine kinase) domain cassette which confers autoregulated kinase activity. Particularly, in these three proteins a carboxy-terminal F (F-actin-binding) domain is identifiable which ties ABL-dependent phosphoregulation to actin filament reorganization (Colicelli, 2010). ABL1 has been implicated to function in a range of cellular processes including actin dynamics and cell migration. ABL was discovered as a cellular proto-oncogene co-opted by the Abelson Murine Leukemia Virus, as well as part of the Breakpoint Cluster Region (BCR)-ABL oncoprotein which is constitutively active in human chronic myelogenous leukemia (CML) and acute lymphocytic leukemia (ALL) (Wang, 2014). Alterations of ABL1 by chromosomal rearrangement or viral transduction can lead to malignant transformation. Activity of the ABL1 protein is negatively regulated by its SH3 domain, the deletion of which turns ABL1 into an oncogene (Barila and Superti-Furga, 1998). Unphosphorylated and inactive murine Abl can be bound by an inhibitor, such as Pag/Msp23, that contacts both the SH3 and the TK domains (Wen and Van Etten, 1997). After removal of the inhibitor, Abl acquires substantial catalytic activity that is further enhanced by primary and secondary (auto)phosphorylation (Brasher and Van Etten, 2000). The ABL kinases have also been secondary (auto)phosphorylation (Brasher and Van Etten, 2000). The ABL kinases have also been shown to play a crucial role in the development of the nervous system. Overexpression of active Abl in adult mouse neurons results in neurodegeneration and neuroinflammation and activation of ABL has been shown to occur in human neurodegenerative disease (Schlatterer et al., 2011). *Drosophila Abl* has a role in axonogenesis and growth cone motility and, of particular relevance here, *Abl* has been implicated in axonal arborization and growth in the *Drosophila* brain (Leyssen et al., 2005).

The adult central brain of *Drosophila* is largely composed of the central complex and the mushroom bodies (MBs). MBs are bilateral and symmetrical structures that are required for learning and memory (Heisenberg, 2003; Busto et al., 2010). Each MB is made of 2000 neurons that arise from 4 neuroblasts whose larval development is known through use of specific markers. Three types of neurons appear sequentially during development: the embryonic/early larval γ, the larval α’ β’ and the late larval/pupal αβ. Each αβ neuron projects an axon that branches to send an α branch dorsally, which contributes to the formation of the α lobe, and a β branch medially, which contributes to the formation of the β lobe (Lee et al., 1999).

APPs have been intensely investigated because of their link to Alzheimer’s disease and neurodegeneration, although the normal *in vivo* function of APPs in the brain remains unclear and controversial. *Drosophila* possesses a single APP homologue, called APPL, expressed in all neurons throughout development. It has been shown that *Appl* is a conserved neuronal modulator of a Wnt planar cell polarity (Wnt/PCP) pathway involved in MB axonal outgrowth (Soldano et al., 2013). It has been proposed that APPL is part of the membrane complex formed by the core PCP receptors. In turn, APPL recruits ABL kinase to the complex and positively modulates DSH phosphorylation (Singh et al., 2010). DSH is an adaptor cytoplasmic protein apparently involved in all known Wnt pathways and, as such, is also a core intracellular component of the Wnt/PCP pathway. It was showed that a 50% reduction of *Abl* leads to a clear enhancement of the *Appl*^*d*^ mutant phenotype. Conversely, wild-type *Abl*^+^ overexpression (*UAS-Abl*) but not *UAS-Abl-KD* (the kinase dead version) within the αβ MB neurons led to a strong rescue of the *Appl*^*d*^ mutant phenotype. Finally, these results, in addition to accompanying biochemical experiments, suggested that APPL regulates the phosphorylation of DSH by ABL kinase and that this mechanism is conserved in mammals (Soldano et al., 2013). Thus, ABL is the key downstream effector of APPL required for MB axon outgrowth.

Huntington’s disease (HD) is a progressive autosomal dominant, neurodegenerative disorder caused by the expansion of a polyglutamine (polyQ) tract at the N-terminus of a large cytoplasmic protein (3144 a.a.), huntingtin (HTT) (Saudou and Humbert, 2016). Several studies indicate that an alteration of wild-type HTT function might also contribute to disease progression (Cattaneo et al., 2005). The fly huntingtin protein (HTT, 3583 a.a.) is similar to the human HTT protein, with four regions of high sequence homology clustered similarly all along the protein in the N-, central- and C-terminal regions. Despite *htt* being highly conserved across all *Drosophila* species, indicating an essential role for biological fitness, null *htt* mutants show no obvious developmental defects (Zhang et al., 2009).

Although ABL is the key downstream effector of APPL required for MB axon outgrowth, its precise role in MB development is poorly documented. Here we show, using analysis of *Abl* loss-of function (LOF) alleles in single-cell MARCM clones that *Abl* is essentially required for axonal growth in the developing αβ neurons of the MBs and that it is expressed in these neurons. Importantly, the overexpression of *Abl* in these neurons leads to similar axonal growth defects to those observed with the LOF allelic combinations, indicating the probable existence of cellular proteins that negatively control neuronal ABL activity. We identify HTT as a potential cellular inhibitor of ABL activity. These results indicate that *Appl* and *htt*, whose human homologs are central players in neurodegeneration, regulate ABL activity in opposite directions to maintain it in the narrow range necessary for normal axon outgrowth.

## RESULTS

### Loss-of-function and gain-of-function of *Abl* induce similar MB αβ neuron phenotype

*Abl* is a component of an APPL Wnt-PCP signaling pathway required for axon growth in MBs (Soldano et al., 2013). Nevertheless, the precise role of *Abl* in MB neurons is poorly described. We have used three *Abl* alleles (*Abl*^*1*^, *Abl*^*2*^ and *Abl*^*4*^) in order to characterize the requirement of *Abl* function in MB development (Fig. S1A). Each of three double heterozygous combinations (*1/2, 2/4* and *1/4*) are adult lethal but are viable at 48 hr after puparium formation (APF) enabling their effects to be investigated at that stage when the αβ neurons are present and can be easily visualized using the anti-Fasciclin II (FASII) antibody (Lee et al., 1999; Reynaud et al., 2015). Interestingly, the three combinations gave similar MB phenotypes with a mixture of wild-type (WT), absence of β lobe, absence of α lobe and absence of α and β lobes (Fig. 1A-D). Control MBs showed less than 0.5% of absence of β lobe (n=402 with 99.5% of the MBs being WT, data not shown). These *Abl* mutant phenotypes, including lethality, were strongly rescued in the presence of an *Abl-GFP* genomic construct (from 20%, n=115 to more than 80%, n=48 of WT MBs p < 10^−5^; Fig. 1D). *Abl* is known to be deleterious when expressed in an unregulated fashion (Schlatterer et al., 2011; Wang, 2014). We overexpressed *Abl* specifically in the MBs using the GAL4/UAS system (Brand and Perrimon, 1993). WT *Abl* overexpression in the MBs under the control of the *OK107-GAL4* line, produced similar phenotypes to those obtained in *Abl* LOF alleles (Fig. 1E; Fig S1B-E). The penetrance of the gain-of-function (GOF) mutant phenotype was even higher than the one obtained with LOF alleles since WT MBs were never detected. Importantly, when a kinase dead version of the *Abl* gene was expressed in the same conditions, no MB mutant phenotype was apparent. This indicates that over-activity of the kinase function of ABL is causing the phenotype. Taken together, these data strongly show that *Abl* expression must be tightly controlled in the MBs in order to ensure normal MB axon development.

**Fig 1.**
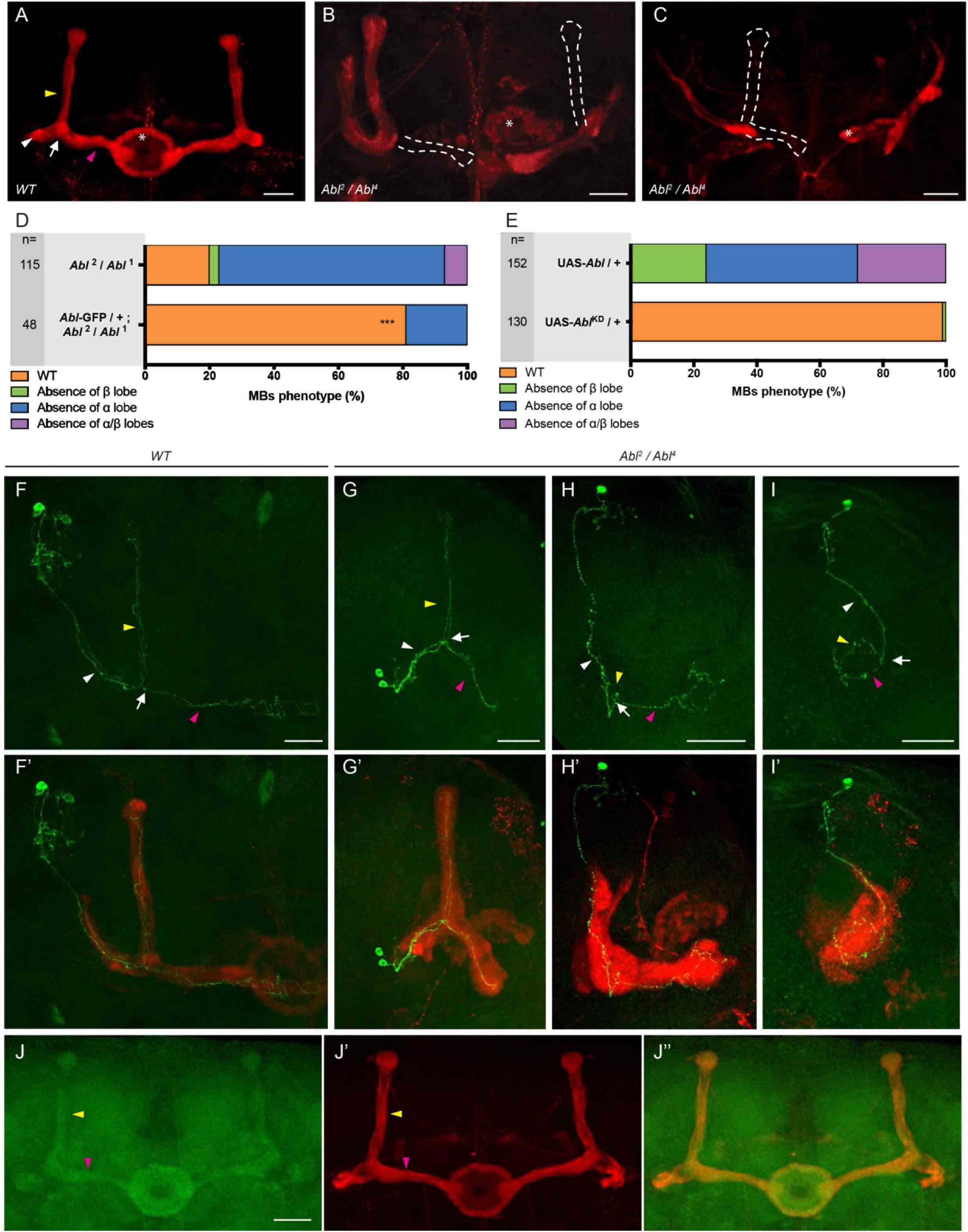
Either loss or overexpression of *Abl* affect MB αβ neuron development. (A-C) Anti-FASII staining of wild-type (WT) brain (A) and on *Abl*^*2*^*/Abl*^*4*^ brain (B-C). In a wild-type (WT) brain, the α lobe (indicated by yellow arrowhead) projects vertically and the β lobe, indicated by pink arrowhead, projects toward the midline and stops before reaching it. The loss of the β and α lobes (B) and of both the α and β lobes (C) is emphasized by white dashed lines. * shows the ellipsoid body. (D) Quantitation of the αβ neuron mutant phenotype in the *Abl*^*2*^*/Abl*^*1*^ mutant and rescued brains with the *Abl-GFP* genomic construct. n= number of MB observed and *** p < 0.001 (Fisher exact test). The *Abl-GFP* genomic construct rescued the developmental defect and lethality of the double heterozygous *Abl*^*2*^/*Abl*^*1*^ combination, but failed to rescue *Abl*^*2*^ homozygote lethality, indicating that this lethality is due to associated modifiers on the chromosome independent of *Abl*. (E) Quantitation of the αβ neuron mutant phenotype when a wild-type or a kinase dead form of *Abl* expression is driven in the MBs by *OK107-GAL4* (n= number of MB observed). (F-F’) Two-cell WT αβ neuron MARCM clone in a WT brain (F) associated with anti-FASII staining in red (F’). (G-G’) Two-cell WT-looking αβ neuron clone (G) associated with anti-FASII staining in red (G’) in an *Abl*^*2*^*/Abl*^*4*^ brain. (H-H’) A single-cell αβ neuron clone with an α branch growth defect (H) associated with anti-FASII staining in red (H’) in an *Abl*^*2*^*/Abl*^*4*^ brain displaying an absence of α lobe. Note the α branch which stops just after the branching point in H (yellow arrowhead). (I-I*’*) A single-cell αβ neuron clone with α and β branch growth and guidance defects (I) associated with anti-FASII staining in red (I’) in an *Abl*^*2*^*/Abl*^*4*^ brain displaying an absence of α and β lobes. Note the small and misguided α and β branches in I. (J-J”) Expression of ABL within the MB using an *Abl-GFP* genomic construct (J). α and β lobes are revealed by anti-FASII staining in red (J’). Merge of GFP and anti-FASII staining (J”). All panels correspond to 48 hr APF brains except for E and the rescue experiment in D which are from adult brains. White arrowheads show the peduncle or common part of the αβ axon, white arrows show the αβ branch point, yellow arrowheads show the α axon branch or the α lobe and pink arrowheads show the β axon branch or the β lobe. The scale bar in panels A-C and F-J indicates 30 µm. Images are composite stacks to allow the visualization of axon trajectories along their entire length. Full genotypes are listed in supplemental information for Fig.1.

### *Abl* function is required for MB axon outgrowth and is expressed in MB αβ neurons

The absence of MB lobes can result from either growth or guidance defects (Soldano et al., 2013; Reynaud et al., 2015). In order to definitively establish which of these two cellular phenomena is affected by loss of *Abl* we produced visualization MARCM clones. *Abl*^*mut*^ two-cell/single cell visualization MARCM clones showed growth defects (87% n = 15) as well as guidance defects (20%) (Table 1 and Fig. 1F-I’). Therefore, we conclude that *Abl*, as *Appl*, is required for MB axon outgrowth. To visualize the localization of ABL protein we employed a genomic *Abl-GFP* construct, which rescues *Abl* mutant lethality and the mutant MB phenotypes (see above) and therefore is likely a bona fide endogenous marker of ABL. *Abl-GFP* is expressed broadly and homogenously in the brain with elevated levels in the MB αβ axons of 48 hours APF brains (Fig. 1J-J”). Taken together, these data show that *Abl* is expressed in the MBs and that *Abl* function is required for MB axon outgrowth.

**Table 1.** Visualization *Abl*^*2*^*/Abl*^*4*^ MARCM clones. WT: wild-type clones. n: number of clones analyzed. Full genotypes are listed in supplemental

| Genotype | WT | $\alpha$ branch growth defect | $\beta$ branch growth defect | $\alpha/\beta$ branches growth and guidance defects | n |
| --- | --- | --- | --- | --- | --- |
| control | 30<br>100% | - | - | - | 30 |
| *Abl<sup>2</sup>/Abl<sup>4</sup>* | 2<br>13% | 10<br>67% | - | 3<br>20% | 15 |

> two-cells clones
| Genotype | WT | Absence of $\alpha$ lobe | Absence of $\beta$ lobe | Absence of $\alpha/\beta$ lobes | n |
| --- | --- | --- | --- | --- | --- |
| control | 9<br>100% | - | - | - | 9 |
| *Abl<sup>2</sup>/Abl<sup>4</sup>* | 6<br>23% | 14<br>54% | 2<br>8% | 4<br>15% | 26 |

Neuroblast clones
| Genotype | WT | Absence of $\alpha$ lobe | Absence of $\beta$ lobe | Absence of $\alpha/\beta$ lobes | n |
| --- | --- | --- | --- | --- | --- |
| control | 9<br>100% | - | - | - | 9 |
| *Abl<sup>2</sup>/Abl<sup>4</sup>* | 2<br>17% | 7<br>58% | - | 3<br>25% | 12 |

### The lack of *htt* rescues MB axon outgrowth phenotypes

Since LOF and GOF of *Abl* produce similar MB phenotypes, its expression must be tightly controlled during normal development. We were therefore interested in isolating potential cellular inhibitors that negatively regulate *Abl* activity in the *Drosophila* MBs. Due to the recessive nature of the *Abl* mutant phenotype and the autosomal location of the gene, we considered looking directly for suppressors to be too laborious. We therefore took advantage of the previous observation that ABL kinase acts downstream of APPL in MB axon growth (Soldano et al., 2013). Flies which are null for *Appl* (*Appl*^*d*^) are viable, fertile and show no gross structural defects in the brain (Luo et al., 1992). The loss of one copy of *Abl* combined with *Appl*^*d*^ (in *Appl*^*d*^ *w\**/Y; *Abl*^*4*^/+ males) increases the MB lobe loss phenotype compared to *Appl*^*d*^ alone, while overexpression of *Abl* in MB neurons rescues the *Appl*^*d*^ phenotype (Soldano et al., 2013). We reasoned that a suppressor of *Abl* activity could also be a suppressor of other components of the Wnt-PCP pathway required for MB axon outgrowth, such as *Appl* and *dsh* for which hemizygous viable mutations are available (*Appl*^*d*^ and *dsh*^*1*^).Therefore, the MB phenotype can be directly assayed in *Appl*^*d*^ males. Fly HTT is a cytoplasmic ubiquitous protein expressed at low levels (Zhang et al., 2009). We identified *htt* as a clear suppressor of the MB *Appl*^*d*^ mutant phenotype. The *Appl*^*d*^ MB phenotype was rescued by reducing *htt* expression using three different genetic manipulations: 1) *htt* heterozygosity using two mutant alleles of *htt* (*htt-ko* or *htt*^*int*^), 2) heterozygosity for a 55 kb chromosomal deficiency uncovering the *htt* locus and most of the adjacent *CG9990*, or 3) *htt* RNAi knock down in the αβ MB neurons (Fig. 2A-E). We noted that loss of one or even two copies of *htt* did not result in any significant MB developmental defects (Fig. S2A). However, we found that reducing one copy of *htt* suppressed also the *dsh*^*1*^ MB phenotype (Fig. 2F-G and J). Hemizygosity for both *Appl* and *dsh* (using a double mutant *Appl*^*d*^ *w dsh*^*1*^ chromosome) resulted in a strong MB axon outgrowth phenotype (up to 60% absence of β lobe, n=129 compared to 21%, n=101 p < 10^−5^ in *dsh*^*1*^ and to 16%, n=141 p < 10^−5^ in *Appl*^*d*^). We found that reducing *htt* was also a strong suppressor of the *Appl*^*d*^ *w dsh*^*1*^ phenotype (12% of absence of β lobe, n=68 p < 10^−5^; Fig. 2 H-I and J). Taken together, these data strongly suggest that HTT is a negative regulator of the Wnt-PCP signaling pathway acting during MB axon outgrowth.

**Fig 2.**
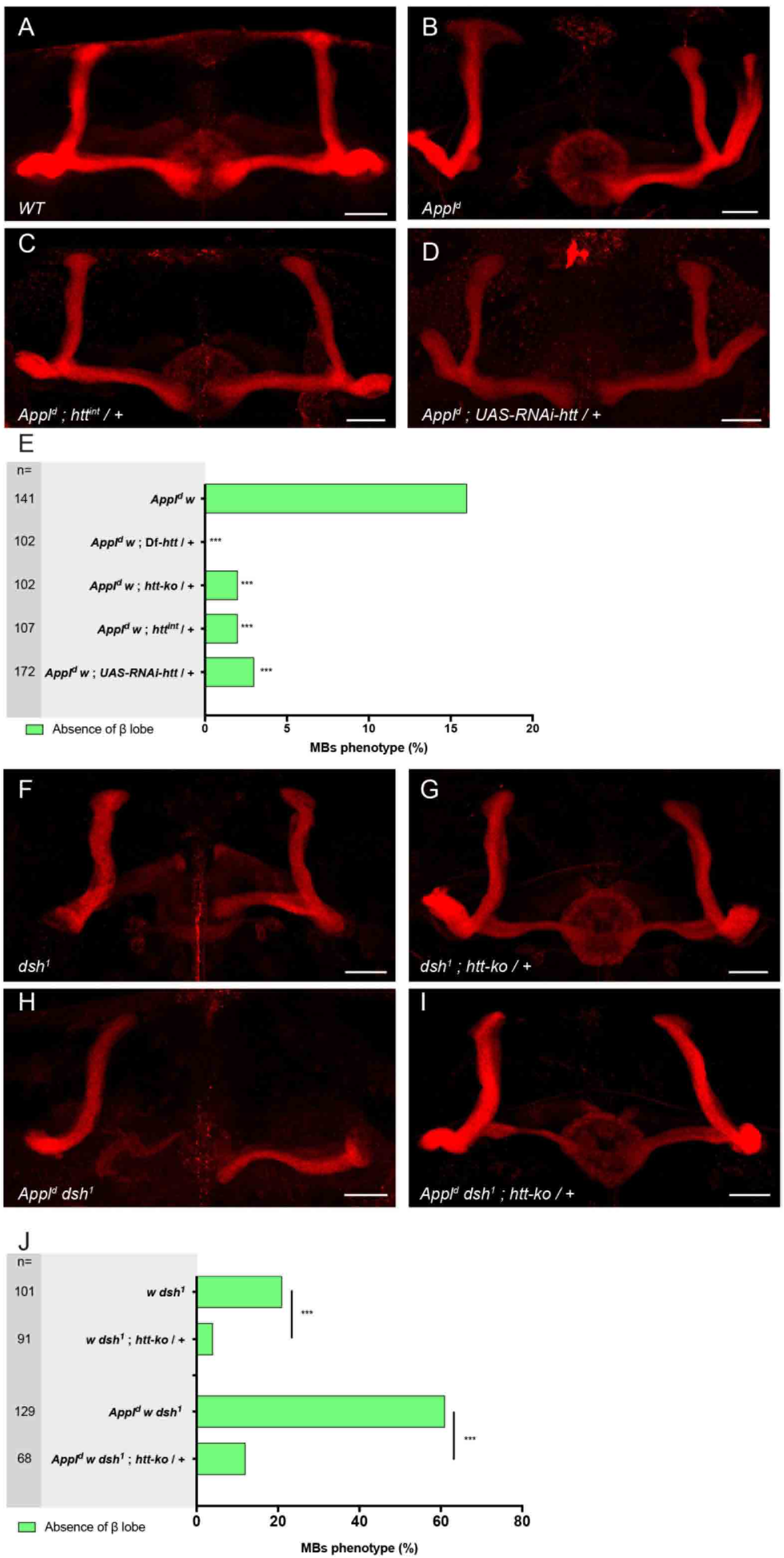
The loss of *htt* rescues the *Appl*^*d*^ and *dsh*^*1*^ MB axon outgrowth mutant phenotype. (A) Wild-type MB α and β lobes revealed by anti-FASII staining. (B-D) Anti-FASII staining reveals the absence of the β lobe in an *Appl*^*d*^ mutant brain (B), which is rescued by the loss of one copy of *htt* (C) and is also rescued by the expression of RNAi against *htt* driven by *c739-GAL4* (D). (E) Quantitation of rescue of the MB *Appl*^*d*^ phenotype by *Df-htt, htt-ko, htt*^*int*^ and *UAS-RNAi-htt* driven by *c739-GAL4*. n = number of MBs analyzed and *** p < 0.001 (Fisher exact test). Note that the rescue of *Appl*^*d*^ by *htt*^*int*^ was done as an independent experiment. Statistics were done from its own *Appl*^*d*^ control, for which the phenotype of β lobe absence was 14% (n = 106; p = 0.0008). (F-G) Anti-FASII staining reveals the absence of β lobe in a *dsh*^*1*^ mutant brain (F), which is rescued by the loss of one copy of *htt* (G). (H-I) Anti-FASII staining reveals the loss of the two β lobes in an *Appl*^*d*^ *dsh*^*1*^ mutant brain (H), which is rescued by the loss of one copy of *htt* (I). (J) Quantitation of the rescue of *dsh*^*1*^ and of *Appl*^*d*^ *dsh*^*1*^ phenotypes by *htt-ko*. n = number of MBs analyzed and *** p < 0.001 (Fisher exact test). All panels correspond to adult brains. The scale bar on panels A-D and F-I indicates 30 µm. Images are composite stacks to allow the visualization of axon trajectories along their entire length. Full genotypes are listed in supplemental information for Fig.2.

### *htt* interacts with *Abl* mutant phenotype in MB axon outgrowth

We found that both the *Appl*^*d*^ and *dsh*^*1*^ MB phenotypes can be rescued by *Abl* expression (Fig. S2B and C). Taken together with the finding that *htt* suppresses both *Appl* and *dsh* MB phenotypes, we hypothesized that *Abl* might be the target of *htt* action. To test this hypothesis we conducted three sets of experiments. First, the increased severity of the MB mutant phenotype in *Appl*^*d*^; *Abl*^*2*^/+ individuals (31% of absence of β lobe, n=102) compared to *Appl*^*d*^ individuals (14% of absence of β lobe, n=100 p=0.0041), was completely abolished when one copy of *htt* was also removed in *Appl*^*d*^; *Abl*^*2*^ *htt*^*int*^/+ individuals (14% absence of β lobe, n=83 with p=0.0089 when compared to *Appl*^*d*^; *Abl*^*2*^/+ individuals and p=1 when compared to *Appl*^*d*^ individuals; Fig. 3A). Second, a modest but significant increase in the proportion of WT MBs in *Abl*^*2*^*/Abl*^*1*^ individuals was observed when one dose of *htt* was also removed (from 20%, n=222 to 32%, n=215 p=0.0086; Fig. 3B; Fig. S3 see discussion). Third, a clear increase of the *Abl* GOF mutant phenotype, measured by the absence of both α and β lobes, was seen when one dose of *htt* was removed (from 21%, n=87 to 75%, n=79 p < 10^−5^; Fig. 3C upper and middle panel). Conversely, a clear rescue of this phenotype was observed when *htt* was also overexpressed using *UAS-htt-fl-CTAP*, a UAS-C-terminally TAP-tagged full length *htt* (from 21%, n=87 to 5%, n=77 p=0.005; Fig. 3C upper and lower panel). Furthermore, the presence of wild-type MBs in these individuals is a strong indication of an interaction between *htt* and *Abl* (from 0%, n=87 compared to 9%, n=77 p=0.0043; Fig. 3C last panel). As expected from these results, expressing *UAS-htt-fl-CTAP* was able to abolish the rescuing effect of *htt*^*int*^/+ on *Appl*^*d*^ males (16% of absence of β lobe in *Appl*^*d*^; *UAS-htt-fl-CTAP/+; htt*^*int*^/+ males, n=120 with p=0.0003 when compared to 2% of absence of β lobe in *Appl*^*d*^; +/+; *htt*^*int*^/+ males, n=107 and p=0.8672 when compared to 17% of absence of β lobe in *Appl*^*d*^ males, n=138) showing its functionality (Fig. 4A). Also, *UAS-htt-fl-CTAP* was properly expressed in the MBs (Fig. 4B). The ability of *htt/*+ to suppress the *Abl* LOF phenotype or to enhance the *Abl* GOF phenotype as well as the ability of *htt* overexpression to suppress the *Abl* GOF phenotype seem to correlate with the amount of kinase-competent residual protein in the various mutant settings (see discussion). Taken together, these data strongly further indicates that HTT is a repressor of ABL function during MB axon outgrowth.

**Fig 3.**
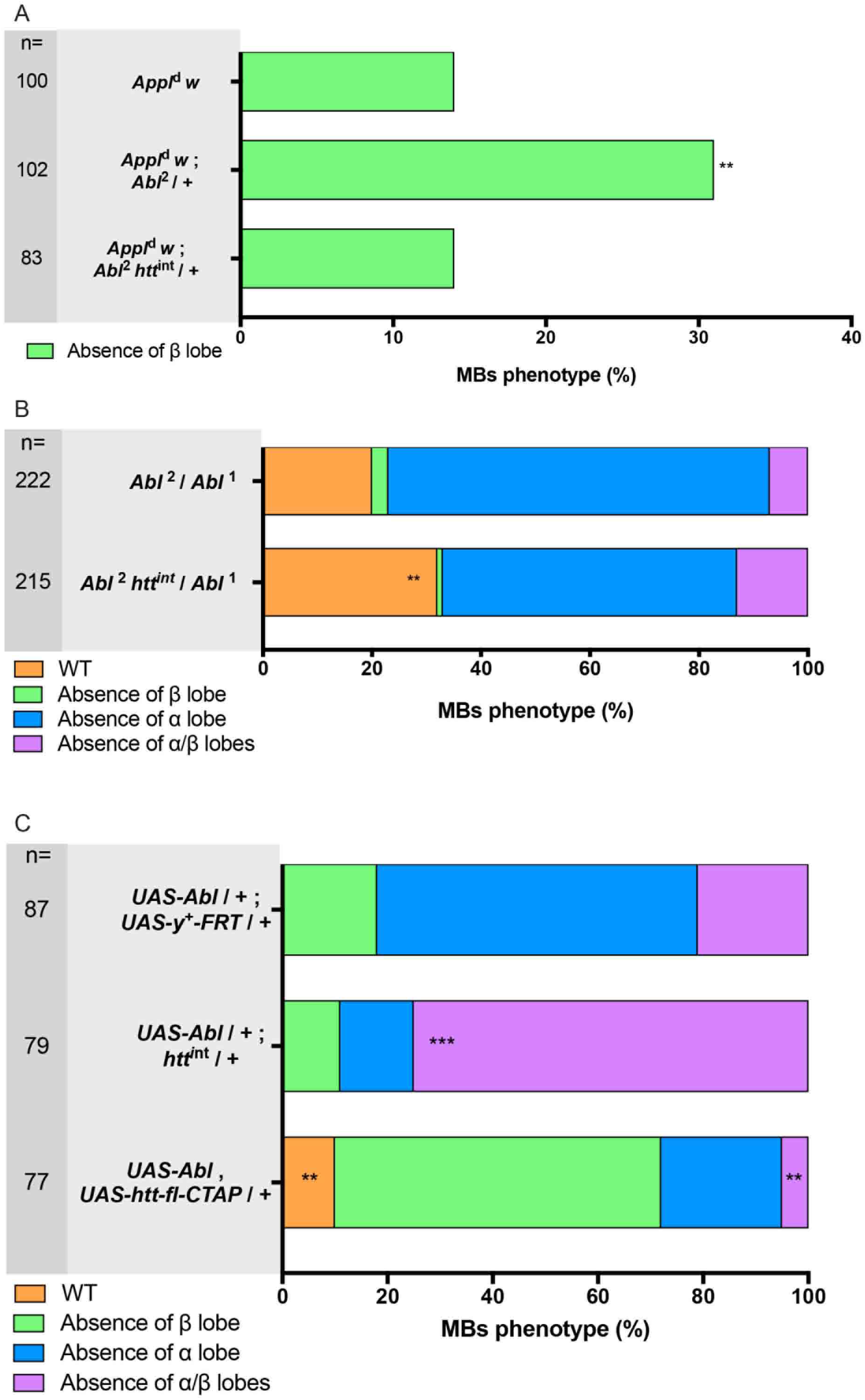
*htt* interacts with *Abl* mutant phenotype. (A) The *Appl*^*d*^ mutant phenotype is enhanced in an *Abl*^*2*^ heterozygote mutant background. However, the *Appl*^*d*^ mutant phenotype is not modified in an *Abl*^*2*^ *htt*^*int*^ double heterozygote mutant background. Note that these two experiments were done independently, but for both of them the *Appl*^*d*^ mutant phenotype was 14% of β lobe absence. (B) The *Abl*^*2*^/*Abl*^*1*^ mutant phenotype is partially rescued by the loss of one copy of *htt* as inferred by the increase of the wild-type (WT) MBs. (C) The absence of α and β lobe phenotype observed in *UAS-Abl* driven by *OK107-GAL4* is strongly increased by the loss of one copy of *htt*. However, this phenotype is rescued by the overexpression of full-length *htt*. The presence of WT MBs in this genotype is also a strong indication of rescue. n = number of MBs analyzed and ** p < 0.01, *** p < 0.001 (Fisher exact test). All panels correspond to adult brains except for B which is from 48 hr APF brains. Full genotypes are listed in supplemental information for Fig.3.

**Fig 4.**
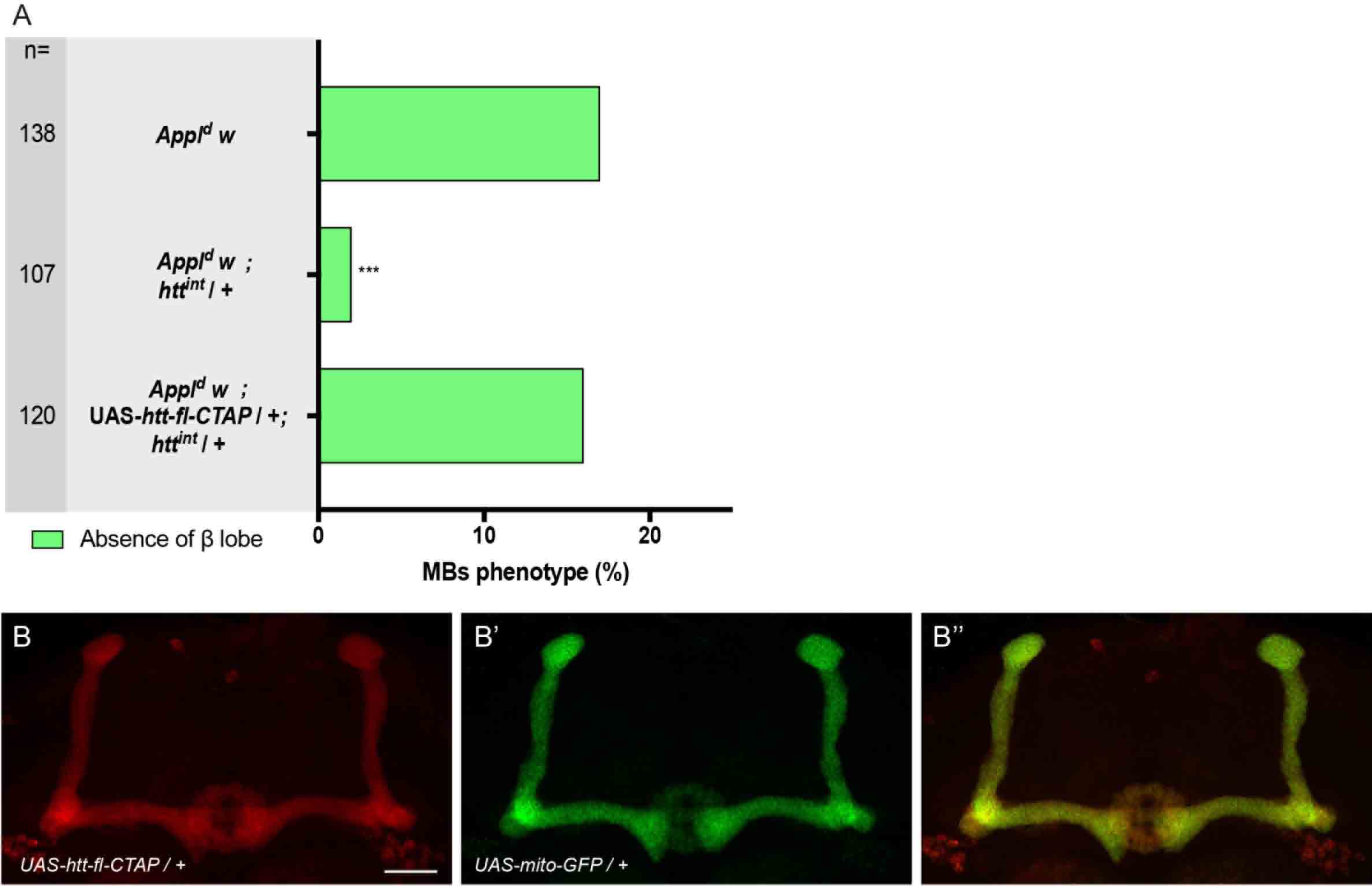
*UAS-htt-fl-CTAP* is functional and properly expressed in the MBs. (A) The rescue of *Appl*^*d*^ by *htt*^*int*^ is prevented by the overexpression of *htt* driven by *c739-GAL4* indicating the functionality of the transgene. n = number of MBs analyzed and *** p < 0.001 (Fisher exact test). Note that *Appl*^*d*^ *w; htt*^*int*^ is also in Fig. 2E. (B-B”) The expression of *UAS-htt-fl-CTAP* revealed by an anti-TAP staining (B) and *UAS-mito-GFP* (B’) driven by *c739-GAL4* are similar in the MBs (B”). All panels correspond to adult brains. The scale bar indicates 30 µm. Images are composite stacks to allow the visualization of axon trajectories along their entire length. Full genotypes are listed in supplemental information for Fig.4.

### Reduction of *htt* increases ABL kinase activity in the developing MBs

One hypothesis to explain the suppressor effect of the lack of one dose of *htt* on the *Abl* MB mutant phenotype is that HTT may have an inhibitory role on ABL kinase activity. A fluorescence resonance energy transfer (FRET) biosensor probe that allows ABL kinase activity to be assayed in living cells was recently introduced and validated in *Drosophila* (Kannan et al., 2017). Using this tool, we first showed that the ABL kinase activity is detectable in the MBs by comparing the FRET activity of the *UAS-Abl*-FRET versus the FRET activity of a mutant that lacks its phosphorylatable tyrosine *UAS-Y→F Abl-*FRET (Fig. 5A-C and E; Fig. S4). We then tested the FRET efficiency in the developing MBs of control (+/+) versus *htt*^*int*^/+ larvae and detected a clear increase of the FRET efficiency when one dose of *htt* was removed (Fig. 5D and F). Quantitative PCR analysis of control versus *htt*^*int*^*/*+ third instar larval brains showed no difference in *Abl* mRNA expression (Fig. 5G). Also, the *htt*/+ heterozygous background does not seem to result in any apparent differences in either the quantity or the localization of ABL protein in the MBs (Fig. 6). Finally, reduced HTT expression does not affect the total levels of neuronal ABL in *Drosophila* heads (Fig. S5). Taken together, these data strongly suggest that HTT is a repressor of ABL kinase activity in the developing MB axons.

**Fig 5.**
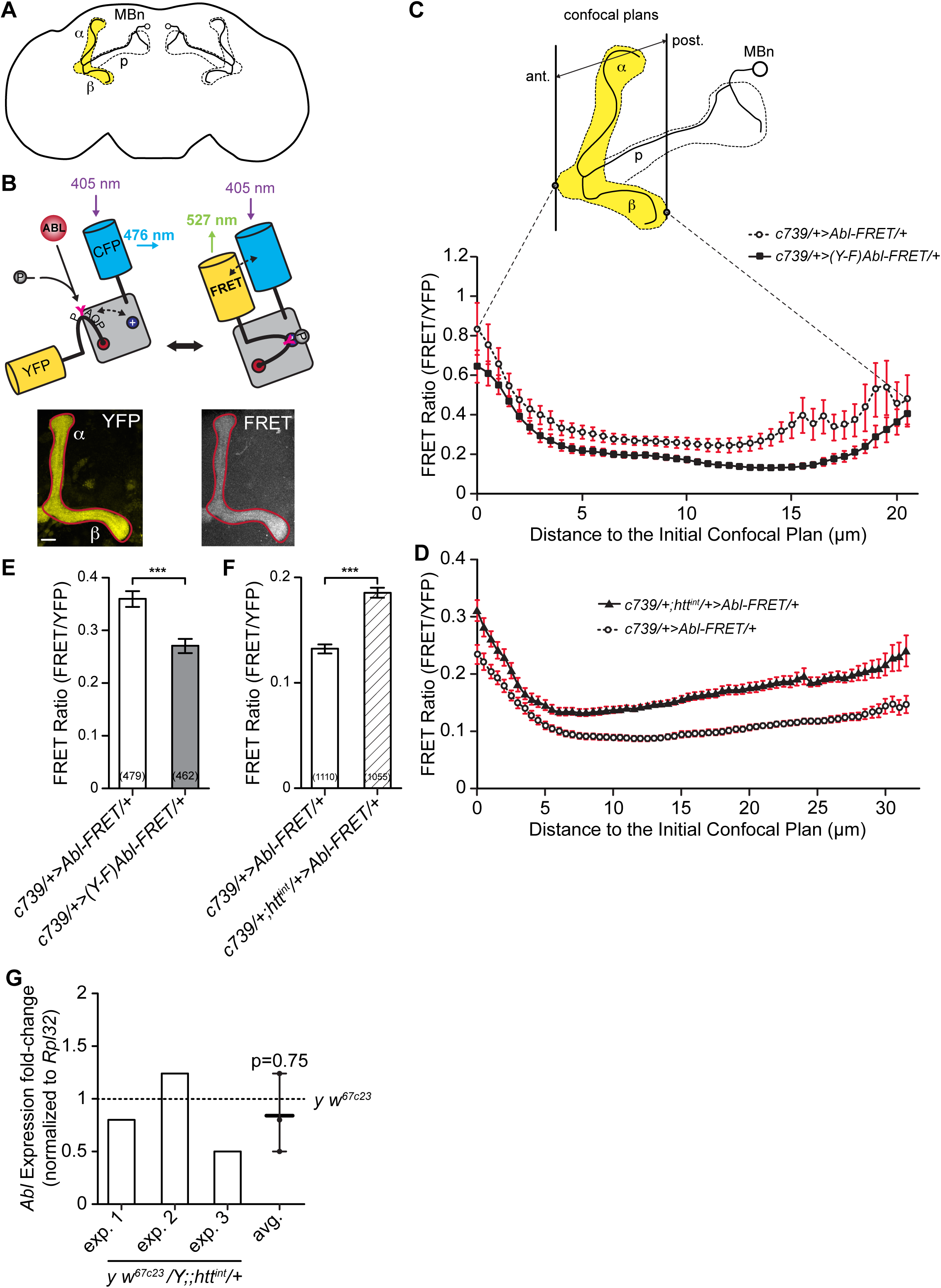
Reducing *htt* expression increases *Abl*-FRET biosensor phosphorylation state during MB development. (A) Schematic representation of an imaged MB in the brain. (B) The *Abl*-FRET biosensor is based on mammalian CRK protein scaffold with two additional fluorescent proteins (CFP and YFP). ABL kinase activity induces phosphorylation of *UAS-Abl*-FRET biosensor leading to its spatial rearrangement and increased FRET efficiency (Ting et al., 2001). Representative maximal projection of the YFP and FRET signals recorded in the MB lobes of adult flies using confocal microscopy. Scale bar: 10µm. (C) FRET and YFP signals are recorded on adjacent 0.5 µm confocal planes along all the anterior-posterior axis of the α and β MB lobes in adult flies. Either wild-type *UAS-Abl*-FRET biosensor or a mutated form (*UAS-Y→F Abl-* FRET) are expressed in the αβ MB neurons using the *c739-GAL4* driver. In the *UAS-Y→F Abl-* FRET biosensor, the tyrosine (Y) located in ABL target site (PYAQP) was replaced by a phenylalanine (F) impairing phosphorylation (Kannan et al., 2017). FRET efficiency is significantly reduced for *UAS-Y→F Abl-*FRET relative to *UAS-Abl*-FRET biosensor for all confocal planes considered except for the nine most anterior and six most posterior planes. Two-tailed Mann-Whitney tests on non-Normally distributed data. Results are mean ± SEM with n ≥ 7 MB for each confocal plane. (D) Reducing *htt* expression increases FRET efficiency of *UAS-Abl*-FRET biosensor in third instar larval MB lobes. The *UAS-Abl*-FRET biosensor is expressed in MB neurons using the *c739-GAL4* driver in control (+/+) and *htt*^*int*^/+ flies. FRET efficiency is significantly increased in *htt*^*int*^/+ flies for all confocal planes along the anterior-posterior axis except for the three located 3 to 4 µm from the initial confocal plane. Two-tailed Mann-Whitney tests on non-Normally distributed data. Results are mean ± SEM with n ≥ 13 MB for each confocal plane. (E) FRET efficiency is globally reduced in *UAS-Y→F Abl-*FRET mutant versus *UAS-Abl*-FRET. FRET efficiency is averaged for all confocal planes and all along the anterior-posterior axis. Two-tailed Mann-Whitney test with non-Normally distributed data. Results are mean ± SEM with n ≥462; *** p<0.001. (F) FRET efficiency is globally increased when *htt* expression is reduced. FRET efficiency is averaged for all confocal planes and all along the anterior-posterior axis. Two-tailed Mann-Whitney test on non-Normally distributed data. Results are mean ± SEM with n ≥ 1055; *** p<0.001. (G) *Abl* mRNA expression is not changed in L3 brains following *htt* partial inactivation. *Abl* expression was assessed using RT-qPCR in L3 brains of *htt*^*int*^ /+ versus WT (+/+) male flies. Results show three independent biological samples. Full genotypes are listed in supplemental information for Fig.5.

**Fig 6.**
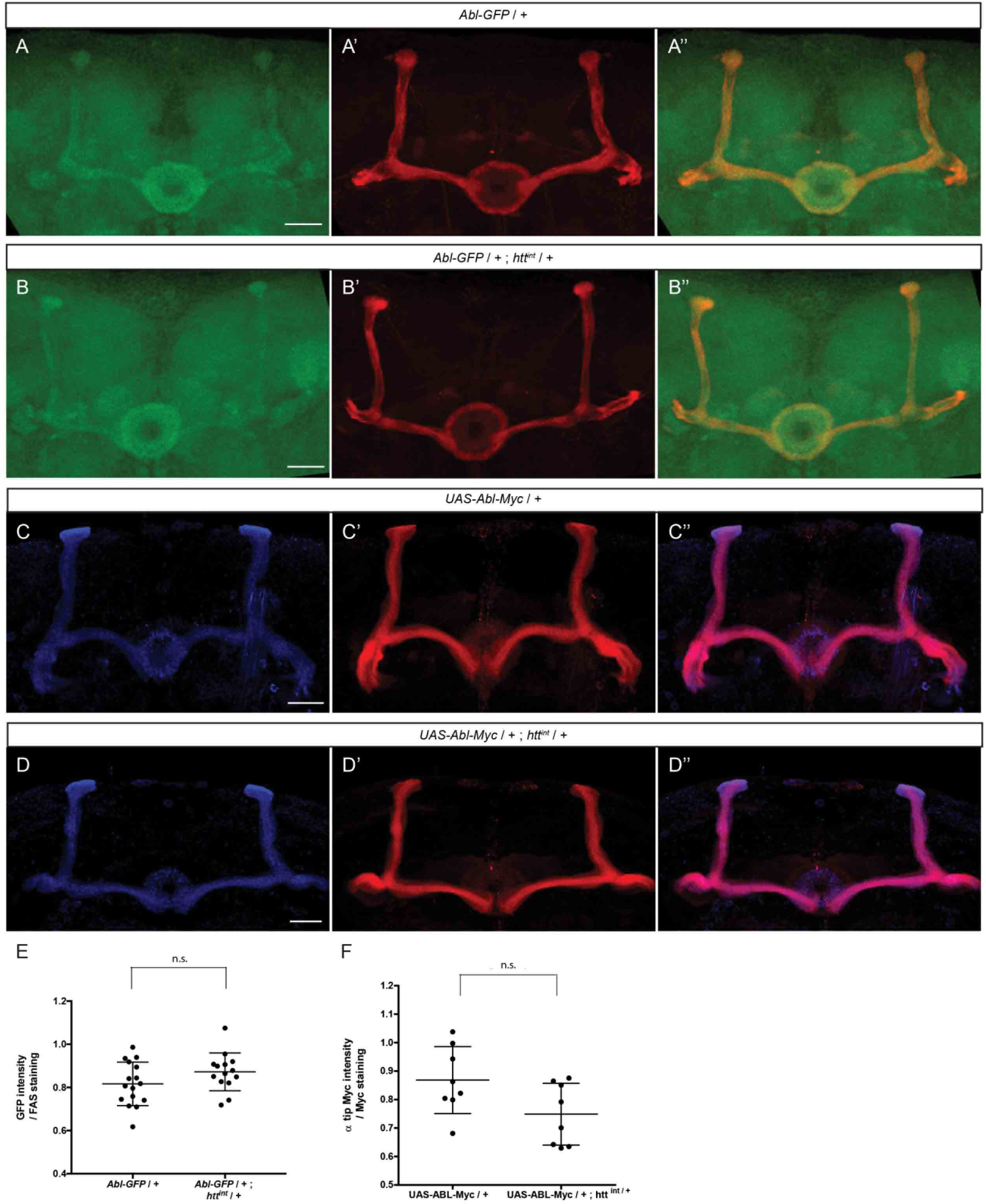
*htt* does not seem to affect the quantity nor the localization of ABL in the MBs. (A-B”) Expression of the genomic construct *Abl-GFP* in a WT (A) and in a *htt*^*int*^ heterozygous mutant background (B) at 48H APF. Anti-FASII staining marked αβ neurons (A’-B’). Merge of GFP and anti-FASII staining (A”-B”). Note that panels A-A” are also presented in Fig. 1 J-J”. (C-D”) Expression of *UAS-Abl-Myc* driven by *c739-GAL4* in a WT (C) and in a *htt*^*int*^ heterozygous mutant background (D) adult brain. Anti-FASII staining marked αβ neurons (C’-D’). Merge of GFP and anti-FASII staining (C”-D”). (E) Quantitation of the GFP expression within MBs is not significantly different between WT and *htt*^*int*^ heterozygous mutant background using a Mann-Whitney *U* test. (F) Quantitation of MYC expression at the tip of the α lobes is not significantly different between WT and *htt*^*int*^ heterozygous mutant background using a Mann-Whitney *U* test. The scale bar indicates 30 µm. Details of image quantification procedure and full genotypes are given in supplemental information for Fig.6.

## DISCUSSION

*Abl* is a key component of the *Appl* signaling pathway required for axonal arborization and growth in the fly brain and the functional relationship between these two proteins is likely conserved in mammals (Leyssen et al., 2005; Soldano et al., 2013). While the role of APP-mediated signaling has been most clearly shown in *Drosophila*, a number of lines of evidence suggest that mammalian APP also fulfills a signaling role (Beckett et al., 2012). Importantly, and in line with its proto-oncogenic role, *Abl* tyrosine kinase activity is tightly regulated by intramolecular inhibition (Barila and Superti-Furga, 1998). Although *Abl* is clearly required as a downstream effector of *Appl* in the MB axon growth, its precise role and regulation in the MBs has not been described.

We show here that *Abl* is required in axonal growth in the MBs, a brain structure that is involved in memory. Furthermore, we show that both *Abl* overexpression and lack of expression in the MBs produce similar phenotypes, indicating the need for tight regulation of ABL activity during MB axon outgrowth. This raises the question of how ABL activity is normally negatively regulated during MB axon outgrowth. We confirmed the previous observation that forced expression of *Abl* rescues the *Appl*^*d*^ MB phenotypes (Soldano et al., 2013). Furthermore, we found that *Abl* overexpression also rescues the *dsh*^*1*^ MB phenotype. These two results support the model that APPL activates ABL, which in turn phosphorylates DSH (Fig. S2B and C). At the genetic level, we expected an increase of ABL activity in an individual bearing LOF allele(s) of a putative *Abl* repressor. We therefore hypothesized that reducing the levels of an *Abl* repressor would result in suppression of the *Appl* and *dsh* mutant MB phenotype. We found that HTT is such a potential inhibitor of ABL activity in the MBs. The loss of one dose of *htt* increased the activity of the wild-type ABL still present in *Abl*^*2*^*/*+ individuals and therefore prevented the enhancement of the MB mutant phenotype (Fig. 3A). While a number of studies have concluded that HTT deficiency results in significant alterations to kinase signaling pathways, to our knowledge this study is the first to implicate the crucial phosphotyrosine kinase, ABL (Bowles and Jones, 2014).

The three *Abl* alleles used in this study are all have been shown to result in truncated proteins (Smith and Liebl, 2005). While *Abl*^*1*^ retains the SH3, SH2 and TK domain of the ABL, *Abl*^*2*^ is mutated within the TK domain and retains the SH3 and SH2 domains and *Abl*^*4*^, which is mutated in the SH2 domain, retains the SH3 domain but completely lacks the TK domain (Fig. S1A). It, therefore, seems likely that very little or no wild type *Abl* function remains in *Abl*^*2*^*/Abl*^*4*^ and *Abl*^*4*^*/Abl*^*1*^ individuals and can possibly explain why no rescue was observed when one dose of *htt* was removed in these genetic backgrounds (Fig. S3). Conversely, since *Abl*^*1*^*/Abl*^*2*^ animals express truncated ABL proteins, this allelic combination is likely less severe than the other two genotypes. Indeed, while significant amounts of truncated *Abl*^*1*^ and *Abl*^*2*^ mutant proteins are detectable, only faint protein bands are observed in *Abl*^*4*^ pupae (Bennett and Hoffmann, 1992). Therefore, some significant kinase activity could remain in *Abl*^*2*^*/Abl*^*1*^ individuals and the loss of one dose of *htt* might increase the activity of the remaining kinase activity (Fig. 3B). Finally, removing one dose of *htt* in animals overexpressing ABL (Fig. 3C middle panel) would release even more ABL activity, which in turn would exacerbate the mutant phenotype. In the opposite way, over-expressing HTT, as in the double *UAS-Abl, UAS-htt* overexpressed individuals (Fig. 3C lower panel), would inhibit ABL function when compared to the *UAS-Abl* overexpression alone which thus might explain the observed rescue.

There are three different levels at which HTT might act to modulate *Abl* activity. First, HTT could either directly or indirectly effect *Abl* mRNA levels. Although fly HTT has been described as a cytoplasmic protein (Zhang et al., 2009), *htt* has been shown to be a suppressor of position-effect variegation, suggesting a role in chromatin organization (Dietz et al., 2015). This hypothesis is unlikely for the MB phenotype described here since qRT-PCR analysis of third instar brains did not reveal a significant difference in *Abl* mRNA levels between *htt*^*int*^/+ and control individuals (Fig. 5G). Second, HTT could play a role in the quantity/stability or the localization of the ABL protein in the MBs. We also consider this unlikely since the quantity of the endogenous protein and the increased localization at the tip of the α lobes observed when *UAS-Abl* is driven by *c739-GAL4* are not different in *htt*^*int*^/+ versus control individuals (Fig. 6). Third, HTT could have an effect on the kinase activity of ABL itself. A FRET biosensor that allowed us to assay ABL kinase activity directly in the MBs showed a clear increase in *htt*^*int*^/+ versus control individuals (Fig. 5). Therefore, we favor the model that HTT acts as an inhibitor of ABL kinase activity during normal development.

The HEAT repeat domains of HTT may function as a solenoid-like structure that acts as a scaffold and mediates inter- and intra-molecular interactions. Other arguments in favor of HTT being a molecular scaffold are its large size and its stability (Saudou and Humbert, 2016). It is tempting to propose that the scaffolding role of HTT in necessary for ABL kinase activity repression. At least in the MBs, a balance between the activity of ABL, positively regulated by the membrane complex formed by the core PCP proteins and APPL, and the repression of ABL activity, which includes HTT might exist. On one hand, if APPL is absent, ABL is not optimally activated leading to defects in MB axon growth. Conversely, when only 50% of HTT is present this leads to de-repression of *Abl* kinase activity, which in turn compensates for its sub-optimal level of activation in the absence of *Appl*. This unexpected apparent balance of activation and inhibition of *Abl* by *Appl* and *htt*, whose orthologs are central players in human disease, may define a conserved functional interaction to maintain ABL activity in the relatively narrow window to appropriately effect axon outgrowth.

## MATERIALS AND METHODS

### *Drosophila* Stocks

All crosses were maintained on standard culture medium at 25°C. Except where otherwise stated, all alleles and transgenes have been described previously (http://flystocks.bio.indiana.edu/). The following alleles were used: *Abl*^*1*^, *Abl*^*2*^, *Abl*^*4*^, *Appl*^*d*^, *dsh*^*1*^, *htt*^*int*^ (Dietz et al., 2015), *htt-ko* and *Df-htt* (Zhang et al., 2009). The following transgenes were used: *UAS-Abl* (#28993), *UAS-Abl*.*K417N* (from #8566) named here *UAS-Abl*^*KD*^ for kinase dead, *UAS-RNAi-htt* (Gunawardena *et* al., 2003), *UAS-RNAi-htt*^*v29351*^ and *UAS-RNAi-htt*^*v29532*^ (VDRC), *UAS-mCD8GFP, UAS-mito-GFP, UAS-FRT-y*^*+*^*-FRT* and the genomic construct *Abl-GFP* (Fox and Peifer, 2007). *UAS-htt-fl-CTAP* was produced for this study (see Constructs). We used three GAL4 lines: *c739-GAL4* expressed in αβ MB neurons, *OK107-GAL4* expressed in all MB neurons (Aso et al., 2009) and the pan-neuronal driver *elav*^*c155*^*-GAL4*. Recombinant chromosomes were obtained by standard genetic procedures and were molecularly checked when required.

### Adult and Pupal Brain Dissection and Immunostaining

Adult brains were dissected in PBS after fly heads and thoraxes had been fixed for 1hr in 3.7% formaldehyde in PBS. They were then treated for immunostaining as previously described (Lee and Luo, 1999; Boulanger et al., 2011). Pupal brains were dissected in PBS and fixed for 20min in 3.7% formaldehyde in PBS at 4 °C with gentle rocking. After washing twice in PBS with 0.5% Triton X-100 (PBT) for 15 min at room temperature, they hey were incubated in PBT and 5% bovine serum albumin (BSA) (blocking solution) at room temperature for 30 min, followed by overnight incubation at 4 °C with primary antibody diluted in blocking solution. Brains were then washed three times in PBS for 20 min, followed by 30 min in the blocking solution, and then addition of the secondary antibody with incubation for 3 hr at 4 °C. Brains were then washed three times in PBS for 20 min and were mounted with Vectashield (Vector Laboratories). Antibody combinations used: anti-Fasciclin II (mAb 1D4 from DSHB) at 1:50 dilution followed by anti-mouse Cy3 (Jackson ImmunoResearch) at 1:300; rabbit anti-MYC (Cell Signaling) at 1:1000 followed by anti-rabbit Cy5 (Jackson ImmunoResearch) at 1:300; mouse anti-TAP (Santa Cruz Biotechnology) at 1:300 followed by anti-mouse Cy3 (Jackson ImmunoResearch) at 1:300.

### MARCM Clonal Analysis

To generate clones in the MB, we used the MARCM technique (Lee and Luo, 1999). For single and two-cell clones, late L3 larvae were heat-shocked at 37°C for 15 min. Brains were dissected at 48 hr APF and then stained. We use the term “visualization MARCM clones” since homozygous mutant clones were examined in a homozygous mutant background.

### Microscopy and Image Processing

Images were acquired at room temperature using a Zeiss LSM 780 laser scanning confocal microscope (MRI Platform, Institute of Human Genetics, Montpellier, France) equipped with a 40x PLAN apochromatic 1,3 oil-immersion differential interference contrast objective lens. The immersion oil used was Immersol 518F. The acquisition software used was Zen 2011. Contrast and relative intensities of the green (GFP), red (Cy3) and blue (Cy5) channels were processed with Fiji Software. Quantitation was performed using ImageJ software.

## Constructs

*pUAS-htt-fl-CTAP*: A full length *htt* “mini-gene” bearing a dual C-terminal Tandem Affinity Purification tag (Protein G and a streptavidin binding peptide: GS-TAP tag) was constructed. PCR was performed using the *dhtt* “mini-gene” comprising the *dhtt* cDNA with intron 10 (as described in Dietz et al., 2015) and the following primers: HTT-TAP-FOR: ggtaccATGGACAAATCCAGGTCCAG (KpnI site added) and HTT-TAP-REV: tctagaCAGGCACTGCAACATCCGG (XbaI site added). The resulting PCR product was digested with KpnI/XbaI and sublconed into the *pUAST-CTAP(SG)* vector (Kyriakakis et al., 2008). To avoid rearrangements due to *dhtt* instability, culturing conditions were used as previously described (Dietz et al., 2015). *pUAS-htt-fl-CTAP* transgenic flies were generated and balanced using standard procedures and expression of dhtt-SG was assessed using western blots.

## FRET Imaging

Fly brains were dissected in 1X PBS at room temperature and collected in ice-cold PBS before being fixed in 3.6% formaldehyde for 20 min. Brains were rinsed twice in PBST 0.5X for 20 min before being mounted in Vectashield® (Vector Laboratories). MBs were imaged using a LSM780 confocal microscope (Zeiss) at x40 with oil immersion and on adjacent 0.5 µm confocal planes along the anterior-posterior axis. Cyan fluorescent protein (CFP) was excited at 405 nm and emission recorded between 454 and 500 nm. Yellow fluorescent protein (YFP) was excited at 514 nm and emission recorded between 516 and 571 nm. Fluorescence resonance energy transfer (FRET) was generated at 405 nm. To avoid CFP emission, FRET was recorded out of CFP emission range, between 587 and 624 nm.

## FRET image analysis

Brains were oriented anterior-posteriorly using the peduncle as an anatomical landmark and aligned according to the first confocal plane where a signal was visible. We ensured that the same number of planes were obtained for each group (*c739>Abl-FRET*: 48 ± 4 planes and *c739>(Y****→****F) Abl-FRET*: 46 ± 1.5 planes, Student t-test: p=0,7. *c739>Abl-FRET*: 79,3 ± 2,3 planes and *c739; htt*^*int*^*>Abl-FRET*: 75,4 ± 3,7 planes; Student t-test: p=0,4) indicating that there were no differences due to mounting. Average YFP and FRET signals were computed using the measurement of ‘Mean Grey Value’ and the ‘Plot Z-axis Profile’ functions of ImageJ (Schneider et al., 2012) into a region of interest (ROI) corresponding to the contour of the MB and for each confocal plane. Background was corrected using the ‘Rolling Ball Background Subtraction’ function (50 px radius). For each plane and within the same ROI, FRET signal was expressed relative to YFP to account for variability in Abl-FRET biosensor expression level or differences between preparations. Only groups (i.e. *Abl-FRET* vs *(Y****→****F) Abl-FRET* and *Abl-FRET* vs *htt*^*int*^*;Abl-FRET*) crossed, collected, dissected and imaged on the same day were compared. For any given confocal plane, FRET ratio was averaged on left and right MBs and multiple animals.

## qRT-PCR

To quantify *Abl* expression, RNA was extracted from the brains of L3 males. Brains (~20/sample) were dissected in PBS 1X (Sigma) and kept on ice before homogenized in TRIzol® reagent (Ambion). Total RNA was treated with DNAse to eliminate genomic DNA (TURBO DNase-*fre*™, Applied Biosystems). RNA was purified using phenol-chloroform extraction and first strand cDNA synthesis was performed using SuperScript^TM^ III reverse transcriptase (Invitrogen). Primers for *Abl* RNA amplification were designed on each side of intron 4-5 within exon 4 and 5 respectively. These exons are present in all *Abl* transcripts. *Abl* primers were designed using Primer3Plus online software and using the qPCR settings (Untergasser et al., 2012). *Abl* forward primer sequence is 5’-GCGGCCATCATGAAGGAAATG-3’ and reverse primer sequence is 5’-TTGCCGTGCGACATAAACTC-3’. *Abl* RNAs were real-time quantified using incorporation of SYBRGreen (Roche) and Light Cycler (Roche). Primers efficacy was first evaluated using a range of cDNA concentrations to ensure linearity of the amplification (E=1,944). Only one melting temperature (Tm=81,6°C) was obtained corresponding to a single PCR product (*Abl* amplicon). The amplicon was run on a gel to ensure the size was the one expected for *Abl* after splicing (107 bp). A control without reverse transcription was done to ensure that *Abl* amplicon was not obtained (Tm≠81.6°C). For each sample, a technical triplicate was performed and averaged. Independent biological replicates were prepared for each condition and the fold change averaged (see Statistics). The biological replicates correspond to independent dissections, extractions, reverse transcriptions and quantifications. In the experimental condition (*y w*^*67c23*^/Y; ; *htt*^*int*^/+), *Abl* expression was expressed relative to control flies (*y w*^*67c23*^/Y) after normalization with internal controls, *Rpl9* and *Rpl32*, and using the ΔΔCT method (Livak and Schmittgen, 2001).

### Western blotting and immunoprecipitations

Lysates of adult *Drosophila* heads were prepared using RIPA buffer supplemented with protease inhibitors (Sigma #11836170001). Antibodies used for immunoprecipitation were: anti-GFP (mouse, HTZ 19C8 and 19F7, Memorial-Sloan Kettering Monoclonal Antibody Facility; (Heiman et al., 2014)), anti-dAbl (rabbit, 1:500; (Song et al., 2010)). Antibodies used for immunoblotting with dilutions were: anti-dhtt (3526, rabbit polyclonal, 1:1,000; (Dietz et al., 2015)), anti-dAbl (as above, 1:1,000), anti-GFP (rabbit, Invitrogen #A-11122, 1:1,000), anti-β-Tubulin (mouse, DSHB E7, 1:10,000), anti-mouse and anti-rabbit IRDye secondary antibodies (LI-COR Biosciences, 1:10,000). Immunoprecipitations were performed in RIPA buffer incubated with 1µg of anti-GFP or 2µg of anti-dAbl. Immunocomplexes were precipitated with either Protein A and Protein G Sepharose beads (anti-GFP), or Protein A Sepharose beads (anti-dAbl) for 1h at 4°C before washing three times with RIPA buffer. Protein A and Protein G Sepharose beads were from Amersham Biosciences. Lysates and immunoprecipitates were resolved on NuPAGE 3-8% gradient Tris-Acetate gels with Tris-Acetate running buffer. After transfer to nitrocellulose membranes, blots were processed according to the Odyssey CLx protocol.

### Statistics

Comparison between two groups expressing a qualitative variable was analyzed for statistical significance using the Fisher exact test (https://www.socscistatistics.com/tests/fisher/Default2.aspx). Comparison of two groups expressing a quantitative variable was analyzed using the two-tailed Mann-Whitney *U* test (https://www.socscistatistics.com/tests/mannwhitney/Default2.aspx). For FRET quantitation, statistical analyses were done using Prism 8.0 (GrapPad). For each confocal plane of each group, normality of FRET ratio was assessed using D’Agostino & Pearson normality test. Non-parametric Mann-Whitney tests were used to compare groups at each confocal plane. For the RT-qPCR, the averaged fold change of *Abl* expression was compared to the theoretical value of 1 that would correspond to no change in *Abl* expression and non-parametric Wilcoxon signed-rank test for non-Normally distributed data or small samples was used. Values of p < 0.05 were considered to be significant.

## Acknowledgments

We thank Florence Maschat and Yoan Arribat for discussions and for providing *htt* fly stocks in an early development of this work, the Bloomington *Drosophila* Stock Center for fly stocks, the imaging facility MRI, member of the national infrastructure France-BioImaging supported by the French National Research Agency (ANR-10-INBS-04, “Investments for the future“)” for the FRET imaging. The 1D4 anti-Fasciclin II hybridoma developed by Corey Goodman and the E7 anti-β-Tubulin monoclonal antibody developed by M. Klymkowsky were obtained from the Developmental Studies Hybridoma Bank, created by the NICHD of the NIH and maintained at The University of Iowa, Department of Biology, Iowa City, IA 52242. C.M. was supported by a PhD grant from the Ministère de l’Enseignement Supérieur et de la Recherche. C.M. and G.B. were supported from the Fondation pour la Recherche Médicale respectively for a 4^th^ PhD year and for a 3 year post-doctoral fellowship. E.G. was supported by funds from the Basic Neuroscience Program of the Intramural Research Program of NINDS, NIH (Z01 NS003013). J.A.W. work was supported by the CHDI Foundation. Work in the laboratory of J.-M.D. was supported by the Centre National de la Recherche Scientifique, the Association pour la Recherche sur le Cancer (grants SFI20121205950 and PJA 20151203422) and the Fondation pour la Recherche Médicale (Programme “EQUIPES FRM2016” project DEQ20160334870).

## Supplemental information

Supplemental information for Fig. 1.

Genotypes: (A) wild type: *y w*^*67c23*^. (B and C): *y w*^*67c23*^; ; *Abl*^*2*^ *FRT2A / Abl*^*4*^ *FRT2A*. (D) top to bottom: *y w*^*67c23*^; ; *Abl*^*2*^ *FRT2A / Abl*^*1*^ *FRT2A. y w*^*67c23*^; *Abl-GFP / +* ; *Abl*^*2*^ *FRT2A / Abl*^*1*^ *FRT2A*. (E) top to bottom: *y w*^*67c23*^ */* Y ; *UAS-Abl / UAS-mCD8-GFP* ; ; *OK107-GAL4 / +*. *y w*^*67c23*^ */* Y; *UAS-Abl*^*KD*^ */ UAS-mCD8-GFP; TM6B,Tb*^*1*^ */ +* ; *OK107-GAL4 /* +. (F) *w*^*\**^ *tubP-GAL80 hs-FLP122 FRT19A / w*^*\**^ *sn FRT19A; c739-GAL4 UAS-mCD8-GFP / UAS-mCD8-GFP. (G-H-I) w*^*\**^ *tubP-GAL80 hs-FLP122 FRT19A / w*^*\**^ *sn FRT19A* ; *c739-GAL4 UAS-mCD8-GFP / UAS-mCD8-GFP* ; *Abl*^*2*^ */ Abl*^*4*^. (J) *y w*^*67c23*^; *Abl-GFP / +*.

Supplemental information for Fig. 2.

Genotypes : (A) *y w*^*67c23*^ / Y ; *c739-GAL4 UAS-mito-GFP / +*. (B) *Appl*^*d*^ *w*^*^/ Y; *c739-GAL4 UAS-mito-GFP / +*. (C) *Appl*^*d*^ *w*^*^ / Y; *c739-GAL4 UAS-mito-GFP /+; htt*^*int*^ */ +*. (D) *Appl*^*d*^ *w*^*^ / Y; *c739-GAL4 UAS-mito-GFP / UAS-RNAi-htt*. (E) top to bottom: *Appl*^*d*^ *w*^*^ / Y; *c739-GAL4 UAS-mito-GFP / +. Appl*^*d*^ *w*^*^ / Y; *c739-GAL4 UAS-mito-GFP / +; Df-htt / +. Appl*^*d*^ *w*^*^ / Y; *c739-GAL4 UAS-mito-GFP / +; htt-ko / +. Appl*^*d*^ *w*^*^ / Y; *c739-GAL4 UAS-mito-GFP / +; htt*^*int*^ */* +. *Appl*^*d*^ *w*^*^ / Y; *c739-GAL4 UAS-mito-GFP / UAS-RNAi-htt*. (F) *w dsh*^*1*^. (G) *w dsh*^*1*^; ; *htt-ko / +*. (H) *Appl*^*d*^ *w*^*\**^ *dsh*^*1*^. (I) *Appl*^*d*^ *w*^*\**^ *dsh*^*1*^; ; *htt-ko / +*. (J) top to bottom: *w dsh*^*1*^. *w dsh*^*1*^; ; *htt-ko / +*. *Appl*^*d*^ *w*^*\**^ *dsh*^*1*^. *Appl*^*d*^ *w*^*\**^ *dsh*^*1*^; ; *htt-ko / +*.

Supplemental information for Fig. 3.

Genotypes: (A) top to bottom: *Appl*^*d*^ *w*^*\**^ / Y; *c739-GAL4 UAS-mito-GFP / +. Appl*^*d*^ *w*^*\**^ / Y; *c739-GAL4 UAS-mito-GFP /* +; *Abl*^*2*^ */ +. Appl*^*d*^ *w*^*\**^ / Y; *c739-GAL4 UAS-mito-GFP / +*; *Abl*^*2*^ *FRT2A htt*^*int*^ */ +*. (B) top to bottom: *y w*^*67c23*^; ; *Abl*^*2*^ *FRT2A / Abl*^*1*^ *FRT2A*. *y w*^*67c23*^; ; *Abl*^*2*^ *FRT2A htt*^*int*^ */ Abl*^*1*^ *FRT2A*. (C) top to bottom: *y w*^*67c23*^ / Y; *UAS-Abl / +; UAS-FRT-y+-FRT / +; OK107-GAL4 / +. y w*^*67c23*^ / Y; *UAS-Abl / +; htt*^*int*^ */ +; OK107-GAL4 / +. y w*^*67c23*^ / Y; *UAS-Abl*, *UAS-htt-fl-CTAP / +* ; ; *OK107-GAL4 /* +. *UAS-FRT-y+-FRT* is used here as a neutral *UAS* to adjust for number of *UAS* sequences.

Supplemental information for Fig. 4.

Genotypes: (A) top to bottom: *Appl*^*d*^ *w*^*\**^ / Y; *c739-GAL4 UAS-mito-GFP / +. Appl*^*d*^ *w*^*\**^ / Y; *c739-GAL4 UAS-mito-GFP / +*; *htt*^*int*^ / +. *Appl*^*d*^ *w*^*\**^ / Y; *c739-GAL4 UAS-mito-GFP / UAS-htt-fl-CTAP*; *htt*^*int*^ / +. (B) *y w*^*67c23*^ / Y; *c739-GAL4 UAS-mito-GFP / UAS-htt-fl-CTAP*.

Supplemental information for Fig. 5.

Genotypes: *y w*^*67c23*^ / Y; *c739-GAL4*/+; *UAS-Abl-FRET/+. y w*^*67c23*^ / Y; *c739-GAL4*/+; *UAS-Y-F Abl-FRET/+. y w*^*67c23*^ / Y; *c739-GAL4*/+; *htt*^*int*^ *UAS-Abl-FRET/+*.

Supplemental information for Fig 6.

(E) After having outlined the MB with the FASII staining, GFP and FASII intensities from MB shape were quantified for each slices of the stack. The GFP intensity of each slices was averaged and then normalized by the mean FASII intensity. Number of MB analyzed: control = 16, *htt* mutant = 14. Quantitation of the GFP expression within MBs is not significantly different between WT and *htt*^*int*^ heterozygous mutant background using a Mann-Whitney *U* test. (F) After having outlined the MB with the FASII staining, MB Myc intensities was quantified from MB shape for each slices of the stack. Specific slices representing MB α tip where defined. α tip Myc intensity was quantified from these specific slices. The α tip Myc intensity of each specific slices was averaged and then normalized by the mean MB Myc intensity. Number of MB analyzed: control = 8, *htt* mutant = 8. Quantitation of MYC expression at the tip of the α lobes is not significantly different between WT and *htt*^*int*^ heterozygous mutant background using a Mann-Whitney *U* test. The scale bar indicates 30 µm. Quantitation were done with ImaJ software. Images are composite stacks to allow the visualization of axon trajectories along their entire length.

Genotypes: (A) *y w*^*67c23*^; *Abl-GFP / +*. (B) *y w*^*67c23*^; *Abl-GFP / +; htt*^*int*^ */ +*. (C) *y w*^*67c23*^; *c739-GAL4 / UAS-Abl-Myc*. (D) *y w*^*67c23*^; *c739-GAL4 / UAS-Abl-Myc; htt*^*int*^ */ +*.

Supplemental information for Table 1.

Genotypes

Control: *w*^*\**^ *tubP-GAL80 hs-FLP122 FRT19A / w*^*\**^ *sn FRT19A; c739-GAL4 UAS-mCD8-GFP / UAS-mCD8-GFP*.

*Abl*^*2*^*/Abl*^*4*^: *w*^*\**^ *tubP-GAL80 hs-FLP122 FRT19A / w*^*\**^ *sn FRT19A; c739-GAL4 UAS-mCD8-GFP / UAS-mCD8-GFP; Abl*^*2*^ */ Abl*^*4*^.

**Fig S1, related to Fig. 1.**
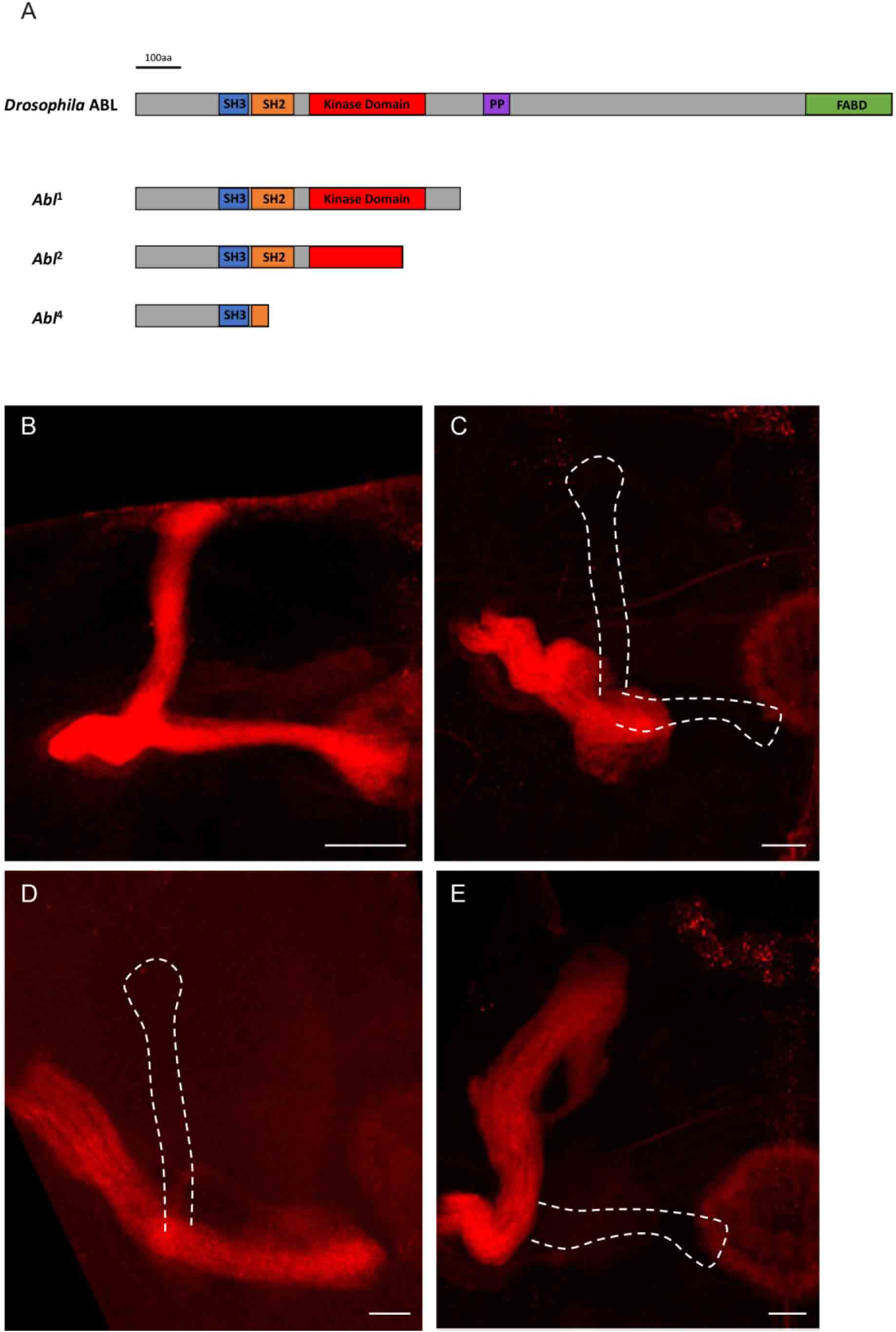
Structure and overexpression of the ABL protein. (A) Molecular scheme of the ABL protein. ABL protein is composed of conserved domains: Src Homology 3 (SH3) domain (blue), Src Homology 2 (SH2) domain (orange), Kinase Domain (red), Poly-Proline PP domain (purple) and F-Actin Binding Domain (FABD) (green). The protein produced by *Abl*^*1*^ mutant allele is truncated between PP and Kinase domains. The protein produced by *Abl*^*2*^ mutant allele is truncated within the Kinase domain. The protein produced by *Abl*^*4*^ mutant allele is truncated within the SH2 domain (Smith and Liebl, 2005). (B-E) Anti-FASII staining showing the α and β lobes in a WT adult brain (B) and in *Abl* forced expression by *OK107-GAL4* (C-E) with an absence of α and β lobes (C), an absence of α lobe (D) and an absence of β lobe (E). Note that panel B is also presented as the left MB in Figure 2A. The scale bar indicates 30 µm. Images are composite stacks. Genotypes: (B) *y w*^*67c23*^ / Y. (C-E) *y w*^*67c23*^ / Y ; *UAS-Abl / UAS-mCD8-GFP* ; ; *OK107-GAL4 /* +

**Fig S2, related to Fig. 2.**
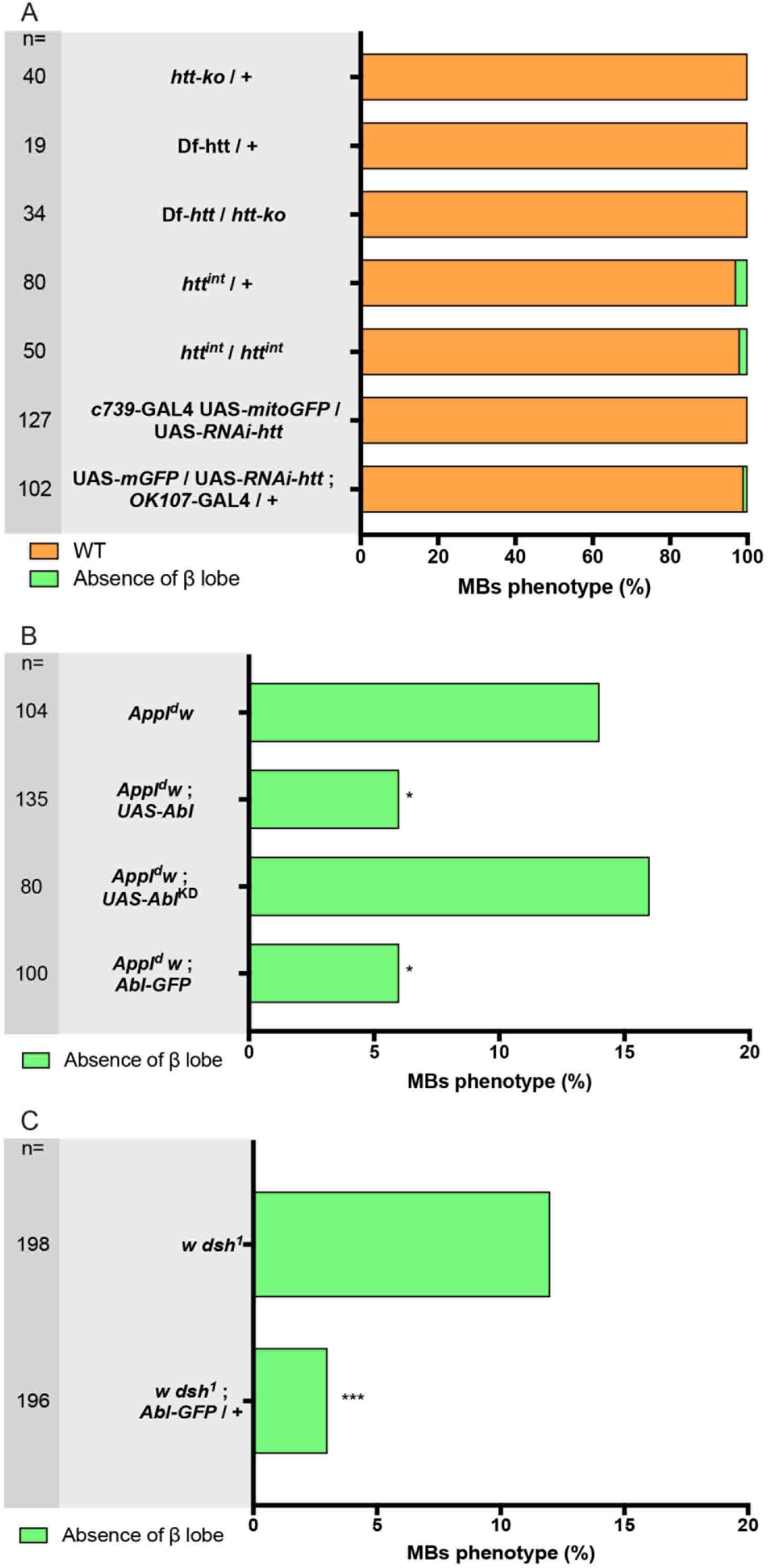
The overexpression of *Abl* rescues the *Appl*^*d*^ and the *dsh*^*1*^ mutant phenotypes. (A) The loss of *htt* does not produce *per se* any significant MB developmental defects. (B) Quantitation of the rescue of *Appl*^*d*^ MB phenotype by *UAS-Abl* driven by *c739-GAL4* and by the genomic construct *ABL-GFP*, but not by the kinase dead form of *Abl, UAS-Abl*^*KD*^. (C) Quantitation of the rescue of *dsh*^*1*^ phenotype by the *Abl-GFP* genomic construct. n = number of MBs analyzed and * p < 0.05, *** p < 0.001 (Fisher exact test). All panels correspond to adult brains. Genotypes: (A) top to bottom: *y w*^*67c23*^ / Y; *c739-GAL4 UAS-mito-GFP / +*; *htt-ko* / +. *y w*^*67c23*^ / Y; *c739-GAL4 UAS-mito-GFP / +*; *Df-htt* / +. *y w*^*67c23*^ / Y; ; *Df-htt / htt-ko*. *y w*^*67c23*^ / Y; ; *htt*^*int*^ */ +*. *y w*^*67c23*^ / Y; ; *htt*^*int*^ */ htt*^*int*^. *y w*^*67c23*^ / Y; *c739-GAL4 UAS-mito-GFP / UAS-RNAi-htt. y w*^*67c23*^/ Y; *UAS-mCD8-GFP / UAS-RNAi-htt* ; ; *OK107-GAL4 / +*. (B) top to bottom: *Appl*^*d*^ *w*^*\**^ / Y ; *c739-GAL4 UAS-mito-GFP / +*. *Appl*^*d*^ *w*^*\**^ / Y; *c739-GAL4 UAS-mito-GFP / UAS-Abl. Appl*^*d*^ *w*^*\**^ / Y; *c739-GAL4 UAS-mito-GFP / UAS-Abl*^*KD*^. *Appl*^*d*^ *w*^*\**^ / Y; *c739-GAL4 UAS-mito-GFP / Abl-GFP*. (C) top to bottom: *w dsh*^*1*^ / Y. *w dsh*^*1*^ / Y; *Abl-GFP* / +.

**Fig S3, related to Fig. 3.**
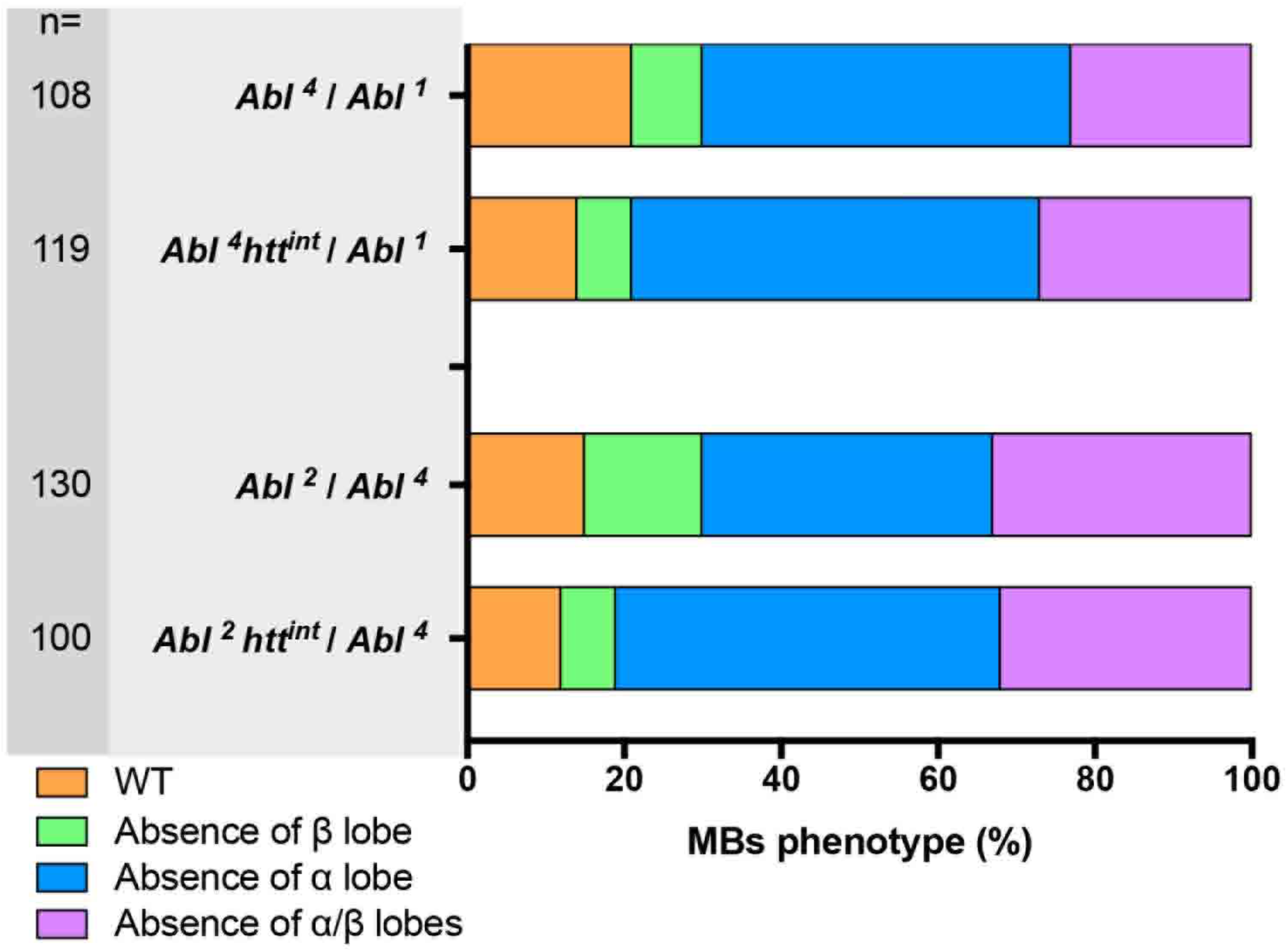
*htt* interaction with *Abl*. The loss of one copy of *htt* does not rescue the *Abl*^*4*^/*Abl*^*1*^ or *Abl*^*2*^/*Abl*^*4*^ mutant phenotype. All panels correspond 48 hr APF brains. Genotypes: top to bottom: *y w*^*67c23*^; ; *Abl*^*4*^ *FRT2A / Abl*^*1*^ *FRT2A*. *y w*^*67c23*^; ; *Abl*^*4*^ *FRT2A htt*^*int*^ */ Abl*^*1*^ *FRT2A*. *y w*^*67c23*^; ; *Abl*^*2*^ *FRT2A / Abl*^*4*^ *FRT2A*. *y w*^*67c23*^; ; *Abl*^*2*^ *FRT2A htt*^*int*^ */ Abl*^*4*^ *FRT2A*.

**Fig S4, related to Fig. 5.**
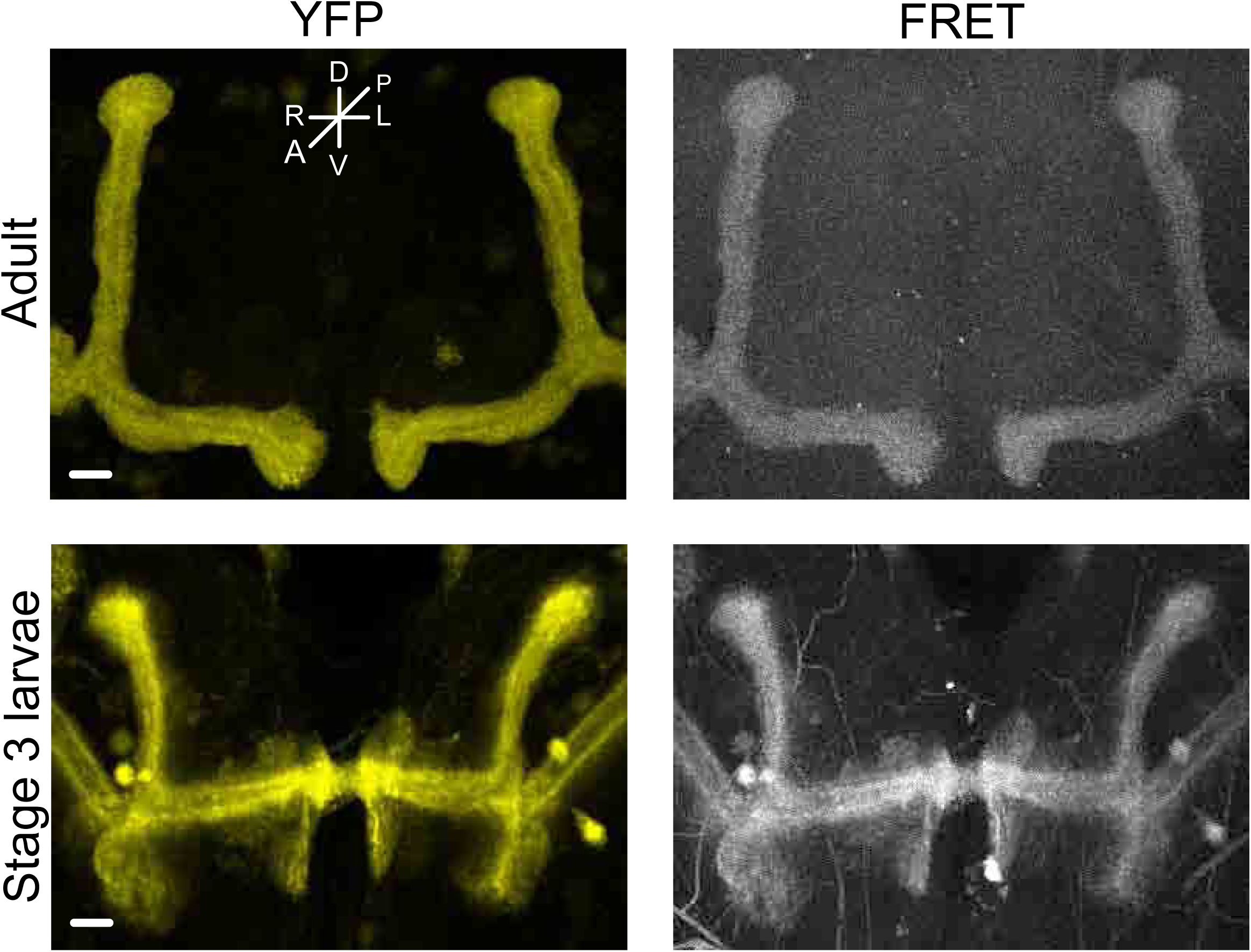
The lack of *htt* releases Abl-FRET biosensor phosphorylation. (*top*) Maximum intensity projection of α and β MB lobes in adult flies. (*bottom*) Maximum intensity projection of vertical and medial MB lobes in stage 3 larvae. The Abl-FRET biosensor is expressed in the MBs using *c739-GAL4* and imaged. Maximum intensity projection of confocal stacks corresponding to YFP and FRET signal are shown. Scale bar: 10µm.

**Fig S5, related to Fig. 6.**
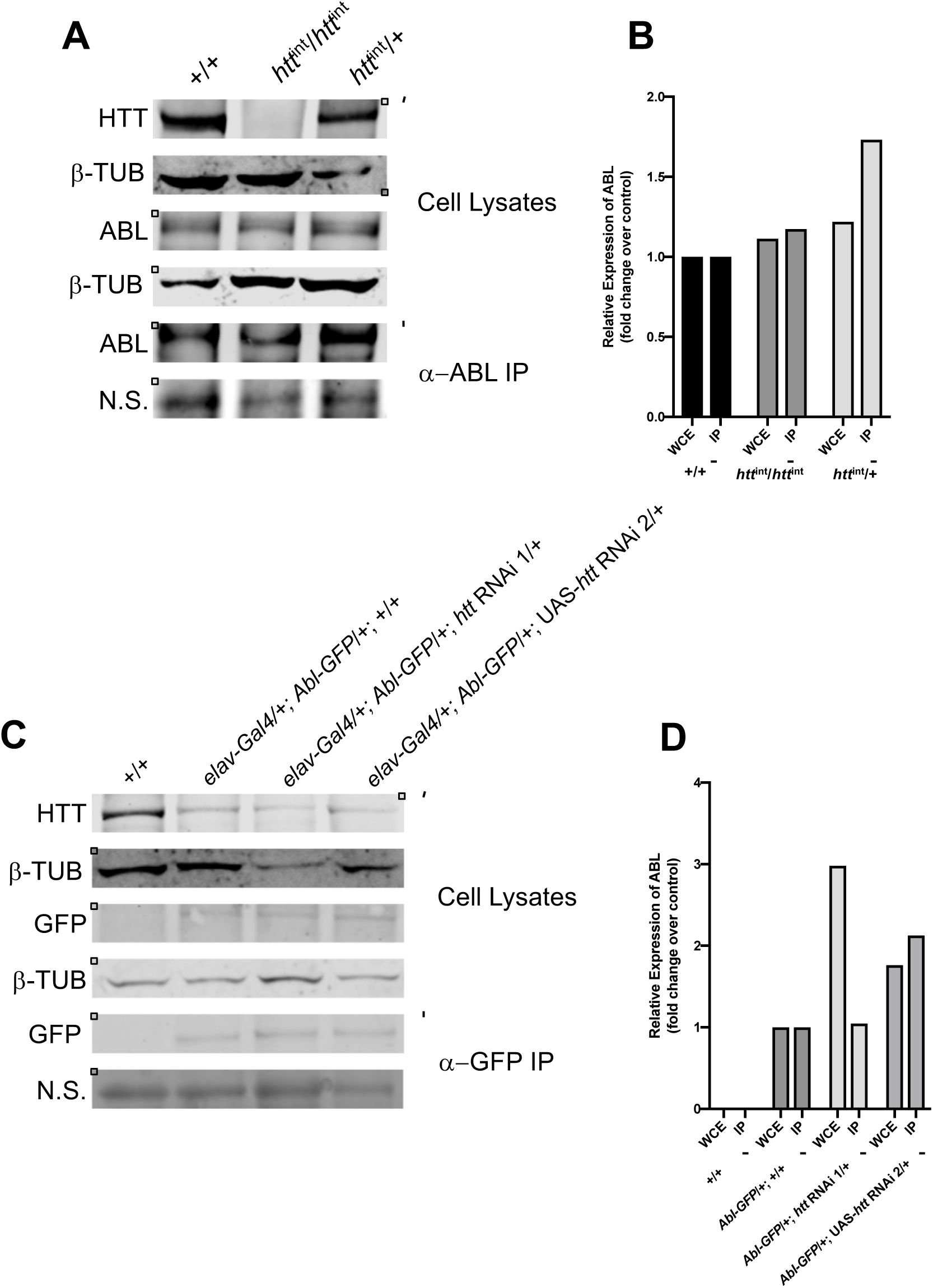
Reduced HTT expression does not affect the total levels of neuronal ABL in *Drosophila*. ABL protein levels are unaffected in either *htt* mutants or with *htt* RNAi knockdown. (A) Whole cell extracts were made from adult heads from the following genotypes: *+/+, htt*^*int*^*/htt*^*int*^ and *htt*^int^/+. Lysates from adult heads were either subjected to western blotting to assess HTT, β-tubulin and ABL levels, or immunoprecipitated using anti-ABL and subjected to western blotting to examine ABL levels. A low molecular weight nonspecific (N.S.) precipitated protein is shown for the IPs. (B) Quantification of ABL protein levels in the indicated genotypes was made from western blots relative to β-tubulin (for cell lysates) or the N.S. band (for anti-ABL IPs) and plotted as fold change in expression relative to the *+/+* control. (C) HTT was knocked down using two RNAi lines (*UAS-htt RNAi*^*v29532*^ and *UAS-htt RNAi*^*v29531*^) driven by *elav* ^*c155*^*-Gal4* in flies expressing GFP-ABL. Lysates from adult heads were either subjected to western blotting to assess HTT, β-tubulin and GFP-ABL (using anti-GFP) levels, or immunoprecipitated using anti-GFP and subjected to western blotting to examine GFP-ABL levels. (D) Quantification of GFP-ABL was assessed relative to β-tubulin (for cell lysates) or N.S.(for anti-ABL IPs) and plotted as fold change in expression relative to the *Abl-GFP* control.

## References

Aso, Y., Grubel, K., Busch, S., Friedrich, A. B., Siwanowicz, I. and Tanimoto, H. (2009) The mushroom body of adult Drosophila characterized by GAL4 drivers. J Neurogenet 23: 156–172.

Barila, D. and Superti-Furga, G. (1998) An intramolecular SH3-domain interaction regulates c-Abl activity. Nat Genet 18: 280–282.

Beckett, C., Nalivaeva, N. N., Belyaev, N. D. and Turner, A. J. (2012) Nuclear signalling by membrane protein intracellular domains: the AICD enigma. Cell Signal 24: 402–409.

Bennett, R. L. and Hoffmann, F. M. (1992) Increased levels of the Drosophila Abelson tyrosine kinase in nerves and muscles: subcellular localization and mutant phenotypes imply a role in cell-cell interactions. Development 116: 953–966.

Boulanger, A., Clouet-Redt, C., Farge, M., Flandre, A., Guignard, T., Fernando, C., Juge, F. and Dura, J. M. (2011) ftz-f1 and Hr39 opposing roles on EcR expression during Drosophila mushroom body neuron remodeling. Nat Neurosci 14: 37–44.

Bowles, K. R. and Jones, L. (2014) Kinase signalling in Huntington’s disease. J Huntingtons Dis 3: 89–123.

Brand, A. H. and Perrimon, N. (1993) Targeted gene expression as a means of altering cell fates and generating dominant phenotypes. Development 118: 401–415.

Brasher, B. B. and Van Etten, R. A. (2000) c-Abl has high intrinsic tyrosine kinase activity that is stimulated by mutation of the Src homology 3 domain and by autophosphorylation at two distinct regulatory tyrosines. J Biol Chem 275: 35631–35637.

Busto, G. U., Cervantes-Sandoval, I. and Davis, R. L. (2010) Olfactory learning in Drosophila. Physiology (Bethesda) 25: 338–346.

Cattaneo, E., Zuccato, C. and Tartari, M. (2005) Normal huntingtin function: an alternative approach to Huntington’s disease. Nat Rev Neurosci 6: 919–930.

Colicelli, J. (2010) ABL tyrosine kinases: Evolution of function, regulation, and specificity. Sci. Signal. 3: re6.

Dietz, K. N., Di Stefano, L., Maher, R. C., Zhu, H., Macdonald, M. E., Gusella, J. F. and Walker, J. A. (2015) The Drosophila Huntington’s disease gene ortholog dhtt influences chromatin regulation during development. Hum Mol Genet 24: 330–345.

Fox, D. T. and Peifer, M. (2007) Abelson kinase (Abl) and RhoGEF2 regulate actin organization during cell constriction in Drosophila. Development 134: 567–578.

Gunawardena, S., Her, L. S., Brusch, R. G., Laymon, R. A., Niesman, I. R., Gordesky-Gold, B., Sintasath, L., Bonini, N. M. and Goldstein, L. S. (2003) Disruption of axonal transport by loss of huntingtin or expression of pathogenic polyQ proteins in Drosophila. Neuron 40: 25–40.

Heiman, M., Kulicke, R., Fenster, R. J., Greengard, P. and Heintz, N. (2014) Cell type-specific mRNA purification by translating ribosome affinity purification (TRAP). Nat Protoc 9: 1282–1291.

Heisenberg, M. (2003) Mushroom body memoir: from maps to models. Nat Rev Neurosci 4: 266–275.

Kannan, R., Song, J. K., Karpova, T., Clarke, A., Shivalkar, M., Wang, B., Kotlyanskaya, L., Kuzina, I., Gu, Q. and Giniger, E. (2017) The Abl pathway bifurcates to balance Enabled and Rac signaling in axon patterning in Drosophila. Development 144: 487–498.

Kyriakakis, P., Tipping, M., Abed, L. and Veraksa, A. (2008) Tandem affinity purification in Drosophila: the advantages of the GS-TAP system. Fly (Austin) 2: 229–235.

Lee, T., Lee, A. and Luo, L. (1999) Development of the Drosophila mushroom bodies: sequential generation of three distinct types of neurons from a neuroblast. Development 126: 4065–4076.

Lee, T. and Luo, L. (1999) Mosaic analysis with a repressible cell marker for studies of gene function in neuronal morphogenesis. Neuron 22: 451–461.

Leyssen, M., Ayaz, D., Hebert, S. S., Reeve, S., De Strooper, B. and Hassan, B. A. (2005) Amyloid precursor protein promotes post-developmental neurite arborization in the Drosophila brain. EMBO J 24: 2944–2955.

Livak, K. J. and Schmittgen, T. D. (2001) Analysis of relative gene expression data using real-time quantitative PCR and the 2(-Delta Delta C(T)) Method. Methods 25: 402–408.

Luo, L., Tully, T. and White, K. (1992) Human amyloid precursor protein ameliorates behavioral deficit of flies deleted for Appl gene. Neuron 9: 595–605.

Reynaud, E., Lahaye, L. L., Boulanger, A., Petrova, I. M., Marquilly, C., Flandre, A., Martianez, T., Privat, M., Noordermeer, J. N., Fradkin, L. G. et al. (2015) Guidance of Drosophila Mushroom Body Axons Depends upon DRL-Wnt Receptor Cleavage in the Brain Dorsomedial Lineage Precursors. Cell Rep 11: 1293–1304.

Saudou, F. and Humbert, S. (2016) The Biology of Huntingtin. Neuron 89: 910–926.

Schlatterer, S. D., Acker, C. M. and Davies, P. (2011) c-Abl in neurodegenerative disease. J Mol Neurosci 45: 445–452.

Schneider, C. A., Rasband, W. S. and Eliceiri, K. W. (2012) NIH Image to ImageJ: 25 years of image analysis. Nat Methods 9: 671–675.

Singh, J., Yanfeng, W. A., Grumolato, L., Aaronson, S. A. and Mlodzik, M. (2010) Abelson family kinases regulate Frizzled planar cell polarity signaling via Dsh phosphorylation. Genes Dev 24: 2157–2168.

Smith, J. A. and Liebl, E. C. (2005) Identification of the molecular lesions in alleles of the Drosophila Abelson tyrosine kinase. Dros. Inf. Serv. 88: 20–22.

Soldano, A., Okray, Z., Janovska, P., Tmejova, K., Reynaud, E., Claeys, A., Yan, J., Atak, Z. K., De Strooper, B., Dura, J. M. et al. (2013) The Drosophila Homologue of the Amyloid Precursor Protein Is a Conserved Modulator of Wnt PCP Signaling. PLoS biology 11: e1001562.

Song, J. K., Kannan, R., Merdes, G., Singh, J., Mlodzik, M. and Giniger, E. (2010) Disabled is a bona fide component of the Abl signaling network. Development 137: 3719–3727.

Staropoli, J. F. (2008) Tumorigenesis and neurodegeneration: two sides of the same coin? Bioessays 30: 719–727.

Thion, M. S. and Humbert, S. (2018) Cancer: From Wild-Type to Mutant Huntingtin. J Huntingtons Dis 7: 201–208.

Ting, A. Y., Kain, K. H., Klemke, R. L. and Tsien, R. Y. (2001) Genetically encoded fluorescent reporters of protein tyrosine kinase activities in living cells. Proc Natl Acad Sci U S A 98: 15003–15008.

Untergasser, A., Cutcutache, I., Koressaar, T., Ye, J., Faircloth, B. C., Remm, M. and Rozen, S. G. (2012) Primer3--new capabilities and interfaces. Nucleic Acids Res 40: e115.

Wang, J. Y. J. (2014) The capable ABL: what is its biological function? Mol. Cell. Biol.

Wen, S. T. and Van Etten, R. A. (1997) The PAG gene product, a stress-induced protein with antioxidant properties, is an Abl SH3-binding protein and a physiological inhibitor of c-Abl tyrosine kinase activity. Genes Dev 11: 2456–2467.

Zhang, S., Feany, M. B., Saraswati, S., Littleton, J. T. and Perrimon, N. (2009) Inactivation of Drosophila Huntingtin affects long-term adult functioning and the pathogenesis of a Huntington’s disease model. Dis Model Mech 2: 247–266.

